# *FUS* gene is dual-coding with both proteins united in FUS-mediated toxicity

**DOI:** 10.1101/848580

**Authors:** Marie A. Brunet, Jean-Francois Jacques, Sonya Nassari, Giulia E. Tyzack, Philip McGoldrick, Lorne Zinman, Steve Jean, Janice Robertson, Rickie Patani, Xavier Roucou

## Abstract

Novel functional coding sequences (altORFs) are camouflaged within annotated ones (CDS) in a different reading frame. We discovered an altORF nested in the FUS CDS encoding a conserved 169 amino acid protein, altFUS. AltFUS is endogenously expressed in human tissues, notably in the motor cortex and motor neurons. Overexpression of wild-type FUS and/or amyotrophic lateral sclerosis-linked FUS mutants is known to trigger toxic mechanisms in different models. These include an inhibition of autophagy, loss of mitochondrial potential, and accumulation of cytoplasmic aggregates. We show here that altFUS, not FUS, is responsible for the inhibition of autophagy. AltFUS is also pivotal in the mechanisms leading to the mitochondrial potential loss and accumulation of cytoplasmic aggregates. Suppression of altFUS expression in a *Drosophila* model of *FUS*-related toxicity protects against neurodegeneration. Some mutations found in ALS patients are overlooked because of their synonymous effect on the FUS protein. Yet we showed they exert a deleterious effect via their missense consequence on the overlapping altFUS protein. These findings demonstrate that *FUS* is a bicistronic gene and suggest that both proteins, FUS and altFUS, cooperate in toxic mechanisms.

## MAIN

FUS is a nuclear RNA-binding protein, with a C-terminal nuclear localization signal (NLS)^1, 2^. The protein is involved in RNA processing, DNA repair and cellular proliferation, although its functions are not precisely elucidated^1^. Mutations in *FUS* gene associate with Amyotrophic Lateral Sclerosis (ALS), frontotemporal lobar dementia (FUS-FTLD) and essential tremor, all characterized by FUS cytoplasmic inclusions in neurons and glial cells^1^. Such cytoplasmic FUS aggregates are pathological features in patients with *FUS* mutations or sporadic disease.

Recently, mutations in *FUS* 3’UTR were described in ALS patients and linked to an increased level of *FUS* mRNA and protein^3–5^. Surprisingly, over-expression of wild-type FUS provokes an aggressive ALS phenotype in mice and fruit flies, in accordance with findings in yeast and mammalian cells^6–10^. The mechanism of the wild-type or ALS-linked mutated FUS toxicity remains unclear^1, 11, 12^.

With currently non-annotated proteins being increasingly reported^13–16^, we hypothesized that the toxicity resulting from wild-type FUS over-expression may come from another, unseen, actor^16^. These novel proteins are coded by alternative open reading frames (altORFs) that are located within “non-coding” RNAs (ncRNA), within the 5’ or 3’ “untranslated” regions (UTR) of mRNAs, or overlapping a known coding sequence (CDS) within a different frame of an mRNA^15–17^. Serendipitous discoveries and ribosome profiling have recently highlighted the distribution of altORFs throughout the human genome, and the consequences of their absence from current databases^16^. For example, mass spectrometry-based proteomics has become the gold standard for protein identification and has been extensively used in ALS studies^18, 19^. However, if a protein is not annotated, it is not included in the protein database (e.g. UniProtKB), and thus cannot be detected by mass spectrometry. An estimated 50 % of mass spectra from a proteomics experiment are unmatched at the end of the analysis^16, 20^.

Genome annotations must avoid spurious ORF annotations. Thus, unless functional characterization has been published, they rely upon 2 arbitrary criteria: a minimum length of 100 codons and a single ORF per transcript. Several groups developed tools to challenge such criteria, such as the sORFs repository^21^ and the OpenProt database^17^, which offer a data-driven broader view of eukaryotic proteomes. The OpenProt database is based on a polycistronic model of ORF annotation^17^ and reports any ORF longer than 30 codons within any frame of an mRNA or ncRNA. It contains currently annotated proteins (RefProts), novel Isoforms, and novel alternative proteins (altProts). Here, we used the OpenProt database (www.openprot.org) to ask whether *FUS* may encode additional proteins that could explain the toxicity of the wild-type protein over-expression. In support of this hypothesis, FUS displays an N-terminal prion-like domain. These low-complexity domains are known to harbour overlapping ORFs^22, 23^.

### ALTFUS IS A NOVEL 170 AMINO ACID PROTEIN, ENDOGENOUSLY EXPRESSED IN CELL LINES AND TISSUES

We began by querying OpenProt^17^ predictions for *FUS* canonical mRNA (*ENST00000254108* or *NM_004960*), which led to 8 predicted altORFs, either overlapping the coding sequence (CDS) or within the 3’UTR (**Appendix Table 1**). Amongst these, IP_243680 or altFUS, a 170 codon altORF overlapping FUS N-terminal prion-like domain, presents convincing experimental evidence of expression (OpenProt v1.3). AltFUS overlaps the FUS CDS in an open reading frame shifted by one nucleotide (**Fig 1A, Fig EV1A**). *FUS* is a complex gene with 13 annotated transcripts resulting from alternative splicing. Based on the GTEx expression data in brain tissues and nerves, five transcripts are more abundant and represent 85 % of all transcripts (**Fig EV1B**). Three of them (*FUS-206*, *FUS-211*, *FUS-203*) are non-coding according to Ensembl, while only two (*FUS-211*, *FUS-203*) are non-coding according to OpenProt (**Fig 1B, Fig EV1C**). Ensembl^24^ annotates two transcripts as coding (*FUS-201* and *FUS-202*), either for the 526 amino acid FUS protein or its 525 amino acid isoform (**Fig EV1D**). From OpenProt prediction, these two transcripts also encode altFUS (IP_243680), or its 169 amino acid isoform (IP_243691) respectively (**Fig EV1E**). Moreover, the second most abundant transcript in brain tissues and nerves (*FUS-206*), representing about 20 % of all transcripts, is non-coding according to Ensembl, but OpenProt predicts it contains the altFUS CDS (**Fig 1B, Fig EV1C**). Thus, of the five most abundant transcripts in brain tissues and nerves, two code for both FUS and altFUS proteins, one codes for altFUS alone, and the remaining two are non-coding.

**Figure 1:**
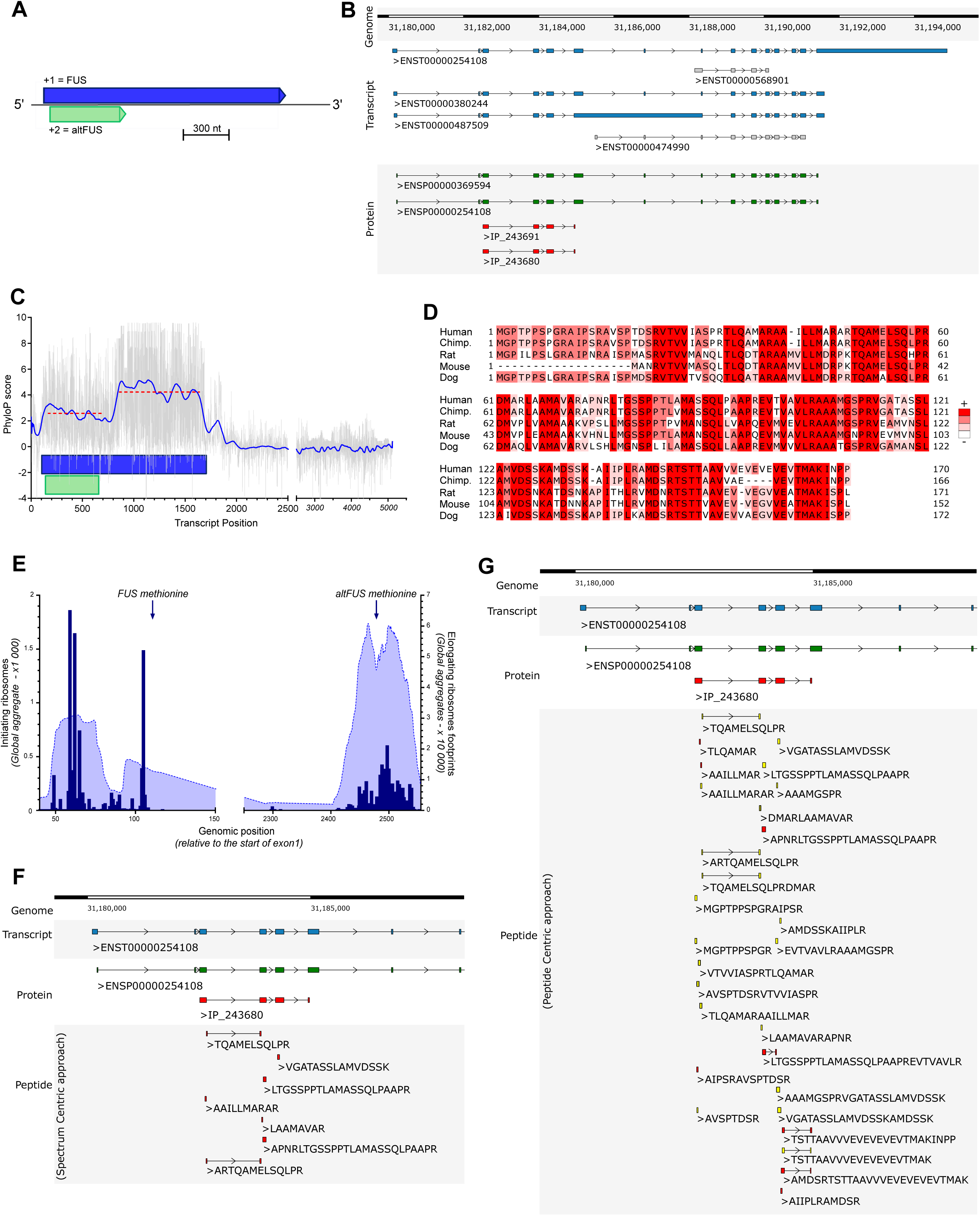
*FUS* is a bicistronic gene **A**, FUS gene bicistronic annotation, with the canonical FUS CDS (in blue, +1 frame) and altFUS CDS (in green, +2 frame) represented on the FUS canonical transcript (*ENST00000254108* or *NM_004960*). Sequence length proportions are respected, the scale bar corresponds to 300 nucleotides. **B**, Genome browser view of FUS gene. The five most abundant transcripts in the brain are shown in the ‘Transcript’ track. Transcripts predicted coding by the OpenProt resource are coloured in blue, and in grey if predicted non-coding. The ‘Protein’ track contains all predicted protein products. The known FUS protein (ENSP00000254108) and its isoform (ENSP00000369594) are coloured in green. The novel predicted altFUS protein (IP_243680) and its isoform (IP_243691) are coloured in red. **C**, PhyloP nucleotidic conservation scores are represented in grey across the *FUS* mRNA (ENST00000254108). The noise reduction after FFT (Fast Fourrier Transformation) is outlined in blue. The average PhyloP score on the bicistronic and the monocistronic region are represented as dotted red lines. The position of the FUS CDS is represented by a blue rectangle, and that of altFUS CDS by a green rectangle. **D**, Alignment (Clustalω) of altFUS protein sequences in Human (*Homo sapiens*), Chimpanzee (Chimp. - *Pan troglodytes*), Rat (*Rattus norvegicus*), Mouse (*Mus musculus*) and Dog (*Canis lupus familiaris*). Residues are coloured based on their identity across species, from white (not conserved) to red (conserved in all species). **E**, RIBO-seq data over the *FUS* gene from the Gwips portal. Initiating ribosome reads are indicated by blue bars, and elongating ribosomes footprints are indicated by the blue curve. The graph capture the beginning of the FUS gene with FUS and altFUS methionines indicated by blue arrows. The genomic positions are indicated relative to the start of exon 1. **F-G**, Genome browser view of *FUS* gene, centered on altFUS. The ‘Transcript’ track contains the beginning of the canonical FUS transcript (*ENST00000254108*) in blue. The ‘Protein’ track contains the beginning of the FUS protein (green) and the whole altFUS protein (red). In **F**, the ‘Peptide’ track contains all the peptides identified by the OpenProt resource using the classical spectrum-centric approach. The peptides sequences are indicated and are unique to altFUS or its isoform (see **Fig EV2B** for an example spectra). In **G**, the ‘Peptide’ track contains all the peptides identified by our peptide-centric approach. The peptides indicated matched better to at least one spectra than any known protein, and are coloured in yellow if they matched better than any known protein with any PTM (see **Fig EV2C** for an example spectra). The peptides sequences are indicated and are unique to altFUS or its isoform.

We retrieved nucleotide conservation scores (PhyloP) for *FUS* transcripts over 100 vertebrates. PhyloP scores range from −10 (highly variable) to 10 (highly conserved). Scores over the FUS CDS are under a constraint at the altFUS CDS locus (average score of 2.6 instead of 4 elsewhere on the FUS CDS), which is consistent with a selection pressure across 2 overlapping frames (**Fig 1C**)^25^. We then retrieved altFUS protein sequences over 84 species and observed a strong protein conservation across mammals, and primates notably (75 to 99.4 % of sequence identity - **Appendix Table 2, Fig EV1F and Appendix Source Data 1**). Thus, AltFUS is well conserved, with domains showing little to no sequence variations (**Fig 1D**). No functional domain nor clear secondary structures could be inferred from bioinformatics predictions (data not shown).

Published RIBO-seq data in Human, retrieved from the Gwips portal^26^, revealed an accumulation of initiating ribosomes around the altFUS initiating methionine, in association with an increase in the density of elongating ribosomes over altFUS CDS (**Fig 1E**). These results suggest that altFUS is translated. Similar results were observed in Mouse (**Fig EV2A**).

Based on the OpenProt database, AltFUS was identified in multiple proteomics datasets, with up to 7 confident peptides (**Fig 1F, Appendix Table 1**). These peptides are unique to altFUS, or shared with its isoform (IP_243691), and represent a 41 % sequence coverage (**Fig EV2B**). Furthermore, we used a peptide-centric approach to query the TCGA datasets for altFUS expression with the PepQuery algorithm. This approach allowed us to identify 28 peptides unique to altFUS, or shared with its isoform (IP_243691), confidently mapped to mass spectra that could not be better explained by any known protein (hg38_Ensembl database) (**Appendix Table 3**). These peptides span through the entire altFUS sequence, representing a full sequence coverage (**Fig EV2C**). Out of these, 20 peptides were confidently mapped to mass spectra that could not be better explained by any known protein with any post-translational modification and/or chemical artefact (**Fig 1G, Fig EV2C**).

To further validate altFUS protein expression, we developed a custom antibody targeting two unique altFUS peptides (**Fig EV3A**) and tested it using three constructs: FUS, altFUS and FUS^(Ø)^. The latter is a monocistronic *FUS* version, where all altFUS methionines are mutated for threonines in a manner synonymous for FUS (**Fig EV3B-D**). Thus, the FUS protein sequence is unchanged, but the altFUS sequence does not contain any methionines. Transfection of HEK293 cells revealed expression of both proteins, FUS and altFUS, from the FUS nucleotide sequence (**Fig 2A**). As expected, altFUS expression was lost with the monocistronic FUS^(Ø)^ construct. HEK293 cells transfected with a siRNA targeting *FUS* mRNA showed a significant knockdown of both proteins, FUS and altFUS; whereas altFUS endogenous expression was visible in scrambled control siRNA and mock transfected cells (**Fig 2B**). These results validate the specificity of the custom antibody for altFUS protein detection by western blot and demonstrate altFUS endogenous expression in HEK293 cultured cells.

**Figure 2:**
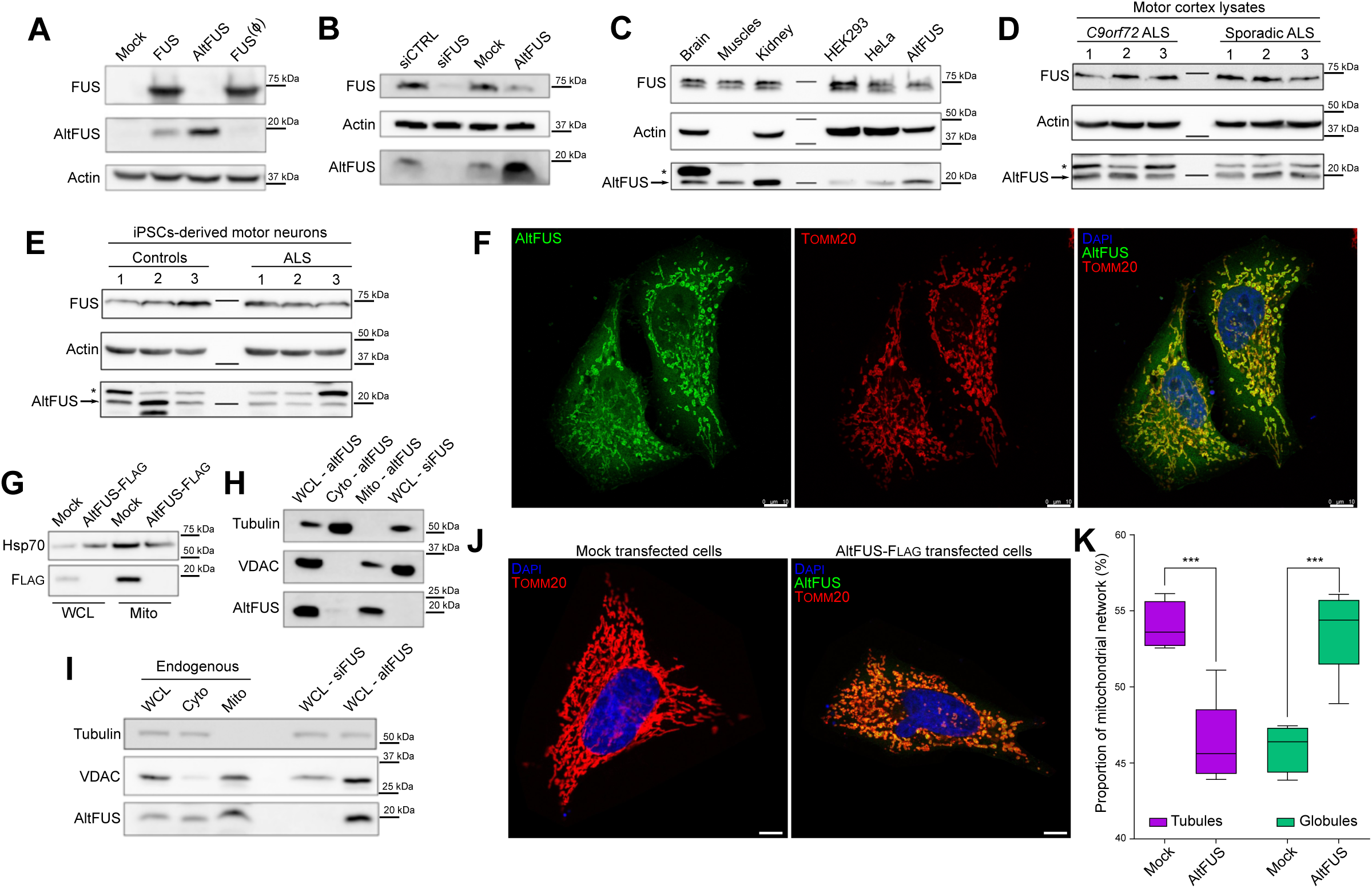
altFUS, a novel endogenous mitochondrial protein. **A**, Expression of both FUS and altFUS proteins from transfection of the *FUS* cDNA in HEK293 cells by western blot, and expression of FUS with the monocistronic construct FUS^(Ø)^ (representative image from n=3). The western blot corresponds to a low exposition so that the endogenous FUS and altFUS are non visible. **B**, AltFUS endogenous expression in HEK293 cells using a siRNA targeting *FUS* mRNA as negative control and over-expression of altFUS CDS as positive control (representative image from n=3). **C**, AltFUS (arrow) endogenous expression in Human tissues (brain, muscles and kidney – 100 ug), in HEK293 and HeLa cultured cells (100 ug) and using the over-expression of altFUS CDS in HEK293 cells (50 ug) as positive control (representative image from n=3). The asterisk indicates a protein species detected with the anti-altFUS antibody specifically in the brain. **D-E**, AltFUS (arrow) endogenous expression in the motor cortex of three *C9orf72* and three sporadic ALS patients (**D**) or in iPSC-derived motor neurons of three lines from controls and from ALS patients (**E**) (representative image from n=3). The asterisk indicates a protein species detected with the anti-altFUS antibody specifically in the brain. **F**, Images by confocal microscopy of altFUS-FLAG (green) in HeLa cells, using TOMM20 (red) as a mitochondrial marker (representative image from n=3, Pearson’s correlation r = 0.92). The white scale bar corresponds to 10 μm. **G**, AltFUS-FLAG enrichment in mitochondrial extracts from transfected HEK293 cells (representative image from n=3) with Hsp70 used as a mitochondrial marker (WCL = Whole Cell Lysate, Mito = Mitochondrial extract). **H**, AltFUS mitochondrial expression in transfected HEK293 cells following fractionation (representative image from n=3), with Tubulin as a cytosolic fraction marker and VDAC as a mitochondrial fraction marker (WCL = Whole Cell Lysate, Cyto = Cytosol fraction, Mito = mitochondrial fraction). **I**, Endogenous altFUS mitochondrial expression in HEK293 cells following fractionation (representative image from n=3), with Tubulin as a cytosolic fraction marker and VDAC as a mitochondrial fraction marker. We used si*FUS* transfected cells as a negative control and altFUS transfected cells as a positive control for altFUS expression (WCL = Whole Cell Lysate, Cyto = Cytosol fraction, Mito = mitochondrial fraction). **J**, Representative images of the mitochondrial network (TOMM20 in red) in mock and altFUS-FLAG (green) transfected HeLa cells (n=3). The white scale bar corresponds to 10μm. **K**, Proportion of tubules and globules in the mitochondrial network of mock and altFUS-FLAG transfected HeLa cells (see **Fig EV4A**). Quantification was done over a minimum of 100 cells across a technical duplicate per independent experiments (n=3, i.e. a minimum of 300 cells per biological conditions, p-value < 0.001, Mann-Whitney U test).

AltFUS endogenous expression was visible in control human tissues, HEK293 and HeLa cell lines (**Fig 2C**). In order to test altFUS expression in pathological tissues, we retrieved motor cortex lysates from 3 ALS patients with a *C9orf72* mutation (most common genetic cause) and 3 sporadic ALS patients (most common aetiology). AltFUS endogenous expression was detected in all cases (**Fig 2D**). Furthermore, as ALS is a motor neuron disease, we derived functional ventral spinal motor neurons from induced pluripotent stem cells (iPSCs)^27, 28^ from healthy controls and ALS patients carrying valosin-containing protein mutations (3 lines per group). AltFUS endogenous expression was detected in all samples (**Fig 2E**). We noticed that brain (**Fig 2C**) and motor cortex lysates (**Fig 2D**), as well as iPSCs-derived motor neurons (**Fig 2E**), from healthy controls and ALS patients, presented a higher band detected with the custom altFUS antibody. This band is not present in cultured cell lines or other tissues. It could come from a non-specific signal or a post-translational modification on altFUS that is specific to the motor cortex and spinal cord motor neurons (**Fig 2C-E**). Deep learning predictions of post-translational modifications on altFUS revealed an extensive propensity to phosphorylation and O-GlcNAcylation (O-linked β-N-acetylglucosamine glycosylation), with up to 19 and 18 sites respectively under a high stringency model (**Appendix Table 4**). These two post-translational modifications are abundant in the eukaryotic brain, and known to be dysregulated in neurodegenerative diseases or aging^29–32^. We could also observe a lower band in the line 2 of controls motor neurons, which may correspond to a degradation product or an initiation at a downstream methionine in altFUS sequence. This band has never been observed in other samples so far. We demonstrate that the *FUS* gene encodes two proteins, FUS and altFUS, both endogenously expressed in human tissues, iPSCs and cell lines.

### ALTFUS IS A MITOCHONDRIAL PROTEIN

FLAG-tagged altFUS (altFUS-FLAG) displayed a strong co-localization with a common mitochondrial marker, TOMM20 (**Fig 2F**). Additionally, mitochondrial extracts showed an enrichment in altFUS-FLAG (**Fig 2G**). Cellular fractionation of cells over-expressing untagged altFUS further validated altFUS mitochondrial localization (**Fig 2H**). The endogenous altFUS protein was found in the mitochondrial fraction, although it displayed a weak cytoplasmic signal as well (**Fig 2I**), consistent with the immunofluorescence data (**Fig 2F**). Furthermore, cells over-expressing altFUS showed an altered mitochondrial network, with a significant increase in fragmented mitochondria (globular) compared to mock cells that displayed more tubular structures (**Fig 2J-K, Fig EV4A**).

Mitochondrial fragmentation is observed in models of overexpression of FUS mutants^33–35^, and we reproduced here a similar effect when over-expressing altFUS alone. Thus, we wondered whether altFUS played a role in other mitochondrial dysfunctions observed in *FUS-*linked toxicity models. To this end, we reproduced an ALS-associated FUS mutant: FUS-R495x^34, 35^. This mutant leads to a premature stop codon before FUS NLS and is linked to severe fALS and sALS cases^1^. In this construct, altFUS is still present and not affected by the mutation (**Fig EV4B**). Similar to FUS^(Ø)^, we also generated the monocistronic construct FUS^(Ø)^-R495x, which contains synonymous mutations for FUS-R495x, but prevents altFUS expression. V5-FUS^(Ø-FLAG)^ and V5-FUS^(Ø-FLAG)^-R495x did not express altFUS, but only the FUS protein, wild-type or ALS-linked mutant R495x respectively (**Fig EV4C**). We first investigated the effect of altFUS on the mitochondrial membrane potential using the potential sensitive dye TMRE (**Fig 3A-B, Fig EV4D**). As previously described^34^, over-expression of bicistronic FUS or FUS-R495x led to a decrease in mitochondrial membrane potential. The mitochondrial membrane potential remained normal when over-expressing monocistronic FUS^(Ø)^ or FUS^(Ø)^-R495x, underlining the role of altFUS. However, over-expression of altFUS alone did not alter the mitochondrial membrane potential, which suggests both proteins cooperate for this FUS-associated toxicity hallmark.

**Figure 3:**
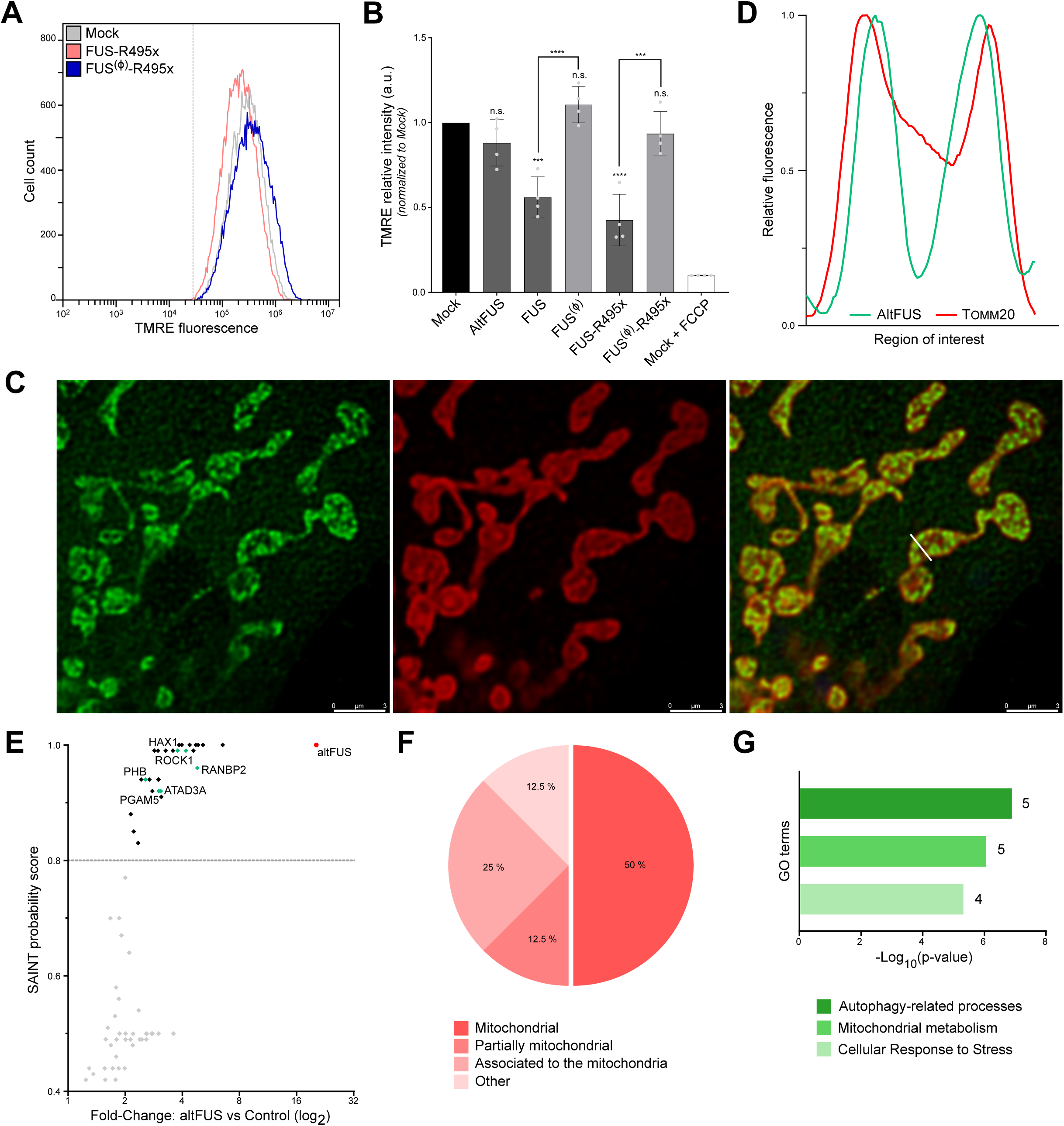
altFUS is involved in mitochondrial dysfunction and autophagy processes. **A**, Representative traces of TMRE fluorescence measured by flow cytometry in mock transfected cells and cells overexpressing the bicistronic FUS-R495x or the monocistronic FUS^(Ø)^-R495x constructs (n=4, minimum of 50 000 live cells per independent replicates). Mean fluorescence intensity of mock transfected cells treated with a decoupling agent, FCCP, is indicated by a grey dotted line. **B**, Mean TMRE fluorescence intensity measures in mock transfected cells, cells over-expressing altFUS, FUS, FUS^(Ø)^, FUS-R495x or FUS^(Ø)^-R495x, or mock transfected cells treated with FCCP across 3 independent experiments (n=4, also see **Fig EV4D**). Statistical significance is relative to the mock condition unless otherwise indicated (*** = p value < 0.001, **** = p value < 0.0001, n.s. = non-significant). **C**, Representative image by stimulated emission depletion microscopy (STED) of altFUS-FLAG (green) localization within mitochondria (TOMM20 marker in red). The white bar across the mitochondria represents the region of interest quantified in panel **D**. The white scale bar corresponds to 3μm. **D**, Relative fluorescence histogram for altFUS-FLAG and TOMM20 across the region of interest highlighted by a white line on panel **C**. **E**, Scatter plot of the proteins identified by AP-MS (see **Fig EV5**) indicating their enrichment (Fold-change over control) and their SAINT probability score. Proteins above the 0.8 threshold are indicated in black, others in grey. AltFUS is indicated in red (bait) and preys known to regulate the autophagy or the cellular stress response are indicated in green. **F**, Subcellular localizations of proteins identified by AP-MS from (see **Fig EV5**). **G**, Enrichment of biological processes in altFUS interacting proteins compared to the Human mitochondrial proteome (n=2, Fisher’s Exact test with FDR < 0.1 %). The number of proteins identified in each GO term is indicated next to the corresponding bar.

To further characterize altFUS, we investigated its protein interactors. Using stimulated emission depletion microscopy (STED), we observed that altFUS localized in puncta following a cristae-like pattern inside the mitochondria, delimited using an outer-membrane mitochondrial marker, TOMM20 (**Fig 3C-D**). We then used size-exclusion chromatography on mitochondrial extracts to isolate altFUS-FLAG macromolecular complexes (**Fig EV5A-B**). Following a FLAG affinity purification and mass spectrometry (AP-MS) analysis, using a no-bait control for quantitative comparison, we confidently identified 12 interacting proteins (**Fig 3E, Fig EV5C-D, Appendix Table 5**). These proteins were identified with a minimum of 2 unique peptides and displayed over a 2-fold enrichment to the control. A gene enrichment analysis to the Human proteome on subcellular localization showed that the identified interactors are in majority proteins known to localize at the mitochondria (**Fig 3F**). Amongst the identified interactors is the heat shock protein HSPA9, a chaperone crucial in the mitochondrial iron-sulfur cluster biogenesis^36^ (**Fig EV5D, Appendix Table 5**). Although HSPA9 was identified with 6 unique peptides and a fold-change of 6.53 to the control, this protein is commonly detected in AP-MS datasets and thus likely non specific nor functionally relevant^37^. Several altFUS interacting proteins are known to interact together, such as PHB, ATAD3A, ERLIN2, RANBP2 and EMD (**Fig EV5D**). A refined gene enrichment analysis to Human mitochondrial proteome identified three significantly enriched biological processes: autophagy-related pathways, mitochondrial metabolism and cellular response to stress (**Fig 3G**). Disruptions within these pathways are pathological hallmarks of *FUS*-linked toxicity and ALS^10, 11, 35, 38^.

### ALTFUS INHIBITS AUTOPHAGY AND DRIVES THE ACCUMULATION OF FUS- AND TDP43-POSITIVE CYTOPLASMIC AGGREGATES

Following on these results, we hypothesized that the inhibition of autophagy observed with ALS-associated FUS mutants may instead be attributed to altFUS. We used the mCherry-GFP-LC3 reporter to track the autophagic flux by confocal microscopy (**Fig EV6A**). Under basal conditions, cells displayed red and yellow foci as expected (**Fig 4A**). An accumulation of yellow foci was observed when cells were treated with bafilomycin, an inhibitor of autophagy. Similarly, cells over-expressing altFUS displayed a significant accumulation of yellow foci (**Fig 4A**). Furthermore, our results were consistent with previously published data^39^ as cells transfected with FUS or FUS-R495x displayed a decreased autophagic flux (**Fig 4A, Fig EV6B**). This accumulation of yellow foci was absent in cells that express monocistronic FUS constructs, thus lacking altFUS expression (FUS^(Ø)^ or FUS^(Ø)^-R495x). We used bafilomycin followed by LC3 probing to further validate the impact of altFUS on autophagy (**Fig 4B, Fig EV6C**). Similarly, an inhibition of autophagy was observed only in cells over-expressing altFUS. Furthermore, in cells over-expressing monocistronic FUS^(Ø)^-R495x, the inhibition of autophagy could be restored by co-transfecting altFUS (**Fig 4B, Fig EV6C**). These results establish altFUS, rather than FUS, as the protein responsible of the inhibition of autophagy.

**Figure 4:**
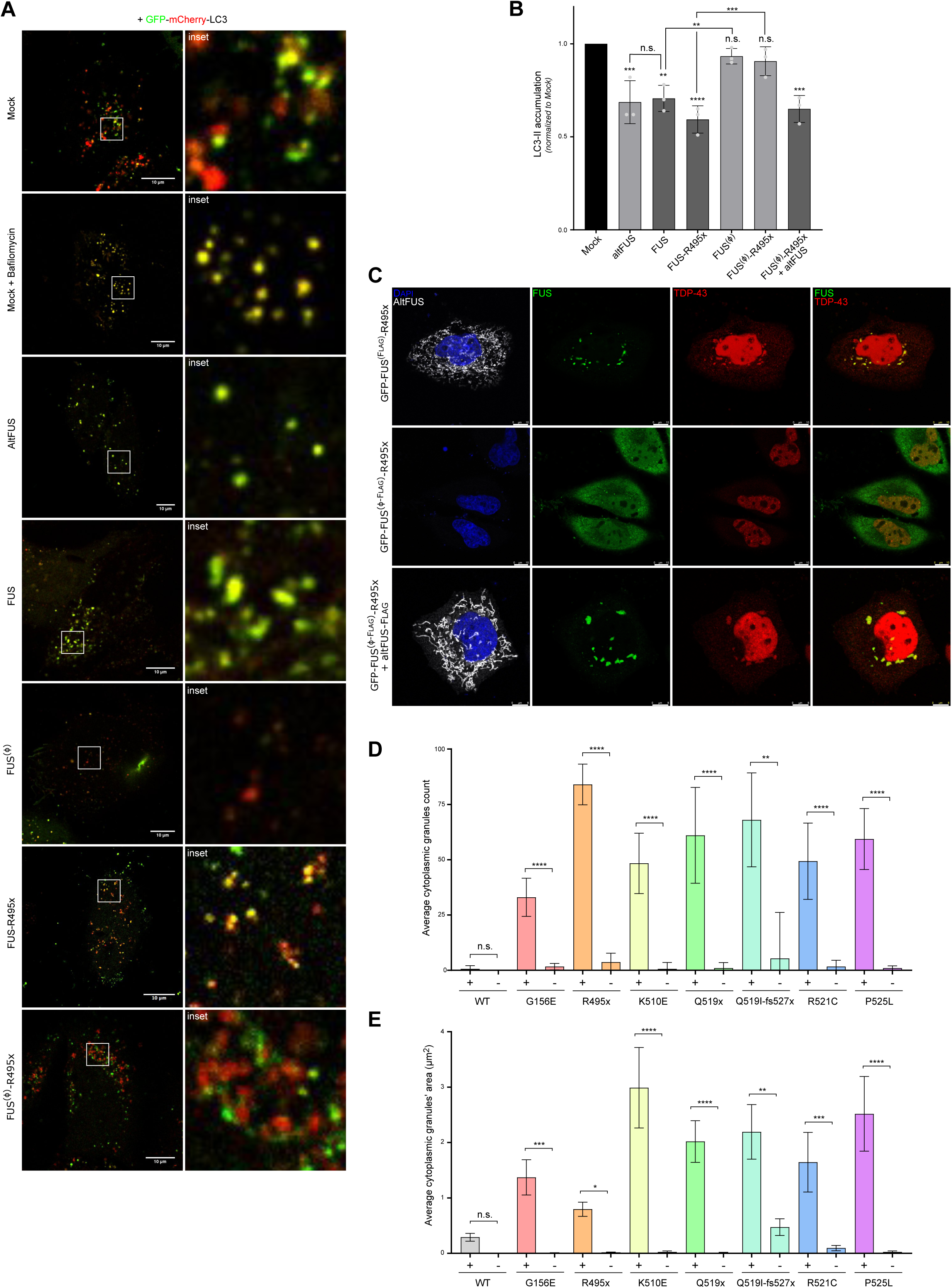
AltFUS is necessary for FUS-associated inhibition of autophagy and accumulation of FUS/TDP-43 cytoplasmic aggregates **A**, Images by confocal microscopy of mCherry-GFP-LC3 signal in HeLa cells across biological conditions: untreated mock, bafilomycin treated mock, altFUS, FUS, FUS^(Ø)^, FUS-R495x and FUS^(Ø)^-R495x (representative images of n=3). The white scale bar corresponds to 10 μm and the zoomed in region (right panels) is delimited by a white box. **B**, LC3-II accumulation after bafilomycin treatment from mock, altFUS, FUS, FUS^(Ø)^, FUS-R495x, FUS^(Ø)^-R495x transfected cells and FUS^(Ø)^-R495x and altFUS co-transfected cells across 3 independent experiments (n=3). The quantification corresponds to the treated/untreated ratio of LC3-II abundance. Statistical significance is relative to the mock condition unless otherwise indicated (**** = p value < 0.0001, *** = p value < 0.001, ** = p value < 0.01, n.s. = non-significant). **C**, Images by confocal microscopy of altFUS (FLAG tagged-white), FUS (GFP tagged - green) and TDP-43 (red) signals in HeLa cells transfected with the bicistronic GFP-FUS^(FLAG)^-R495x or the monocistronic GFP-FUS^(Ø-FLAG)^-R4955x constructs, or co-transfected with the monocistronic GFP-FUS^(Ø-FLAG)^-R495x and altFUS-FLAG constructs (representative images from n=3). The white scale bar corresponds to 10 μm. **D-E**, Quantification of FUS cytoplasmic granules, number (**D**) and area (μm^2^) (**E**) in cells over-expressing the bicistronic (+) or monocistronic (-) construct for FUS, FUS-G156E, FUS-R495x, FUS-K510E, FUS-Q519x, FUS-Q519I-fs527x, FUS-R521C, and FUS-P525L. Statistical comparisons are made between bicistronic and monocistronic versions of each construct (**** = p value < 0.0001, *** = p value < 0.001, ** = p value < 0.01, * = p value < 0.05, n.s. = non-significant, one-way ANOVA test with Sidak’s multiple comparison).

Furthermore, altFUS interactome analysis suggested a role in the cellular stress response, which is known to be altered in ALS with a TDP-43 cytoplasmic accumulation in 98 % of patients^40, 41^. In *FUS*-linked ALS and some sALS cases, FUS cytoplasmic aggregates or mislocalization are also observed^42–44^. In our hands, cells over-expressing FUS-R495x displayed cytoplasmic aggregates that were positive for both FUS-R495x and TDP-43 (**Fig 4C**). Although TDP-43 aggregates are not a common observation with FUS-associated mutants, co-aggregation has already been reported in patients, animal models and cultured cell lines across multiple studies^42, 44–49^. In cells over-expressing the monocistronic FUS^(Ø)^-R495x construct, thus lacking altFUS expression, FUS-R495x displayed a more diffuse cytoplasmic localization, and TDP-43 remained in the nucleus (**Fig 4C**). FUS cytoplasmic aggregates were significantly more numerous and larger when altFUS was co-expressed (**Fig 4D-E**). Accumulation of FUS-R495x and TDP-43 in cytoplasmic aggregates could be reconstituted by co-transfecting altFUS and the monocistronic FUS^(Ø)^-R495x construct (**Fig 4C**). These observations were repeated across all 7 ALS-associated FUS mutations tested (**Fig EV6D-F**). The cytoplasmic aggregates were also TIAR-positive as observed with ALS-linked FUS mutants in previous work^40^ (**Fig EV7**). These results suggest that altFUS enhances the assembly of cytoplasmic FUS mutants aggregates and is responsible for the recruitment of TDP-43 in these aggregates.

### ALTFUS PROTECTS AGAINST NEURODEGENERATION IN FUS-ASSOCIATED DROSOPHILA MODELS

In order to investigate the role of altFUS in an already established *in vivo* model of FUS-related neurodegeneration, we generated *Drosophila* models expressing either the bicistronic, FUS and FUS-R495x constructs, or the monocistronic, FUS^(Ø)^ and FUS^(Ø)^-R495x, constructs. We used the Elav-GeneSwitch-GAL4 Driver strain, as previously described^50^, as it allows for an inducible over-expression in motor neurons and avoids lethality at the larval stage from FUS over-expression in the central nervous system^50, 51^. First, we generated flies containing the sequences for UASt-altFUS, UASt-FUS, UASt-FUS^(Ø)^, UASt-FUS-R495x or UASt-FUS^(Ø)^-R495x. These flies were then crossed with the Elav-GeneSwitch-GAL4 driver strain (**Fig 5A**). UASt-mCherry flies were used as controls. Selected F1 individuals were then divided into 2 groups with equal proportions of males/females. The first group received standard food, while the other received RU-486 treated food. The treatment induces a conformational change in the Elav-GeneSwitch driver, which allows activation of the UAS promoter and thus expression of the target protein. We retrieved flies at selected time points to validate protein expression in the RU-486 treated population through time, while the controls showed no expression (**Fig 5B**).

**Figure 5:**
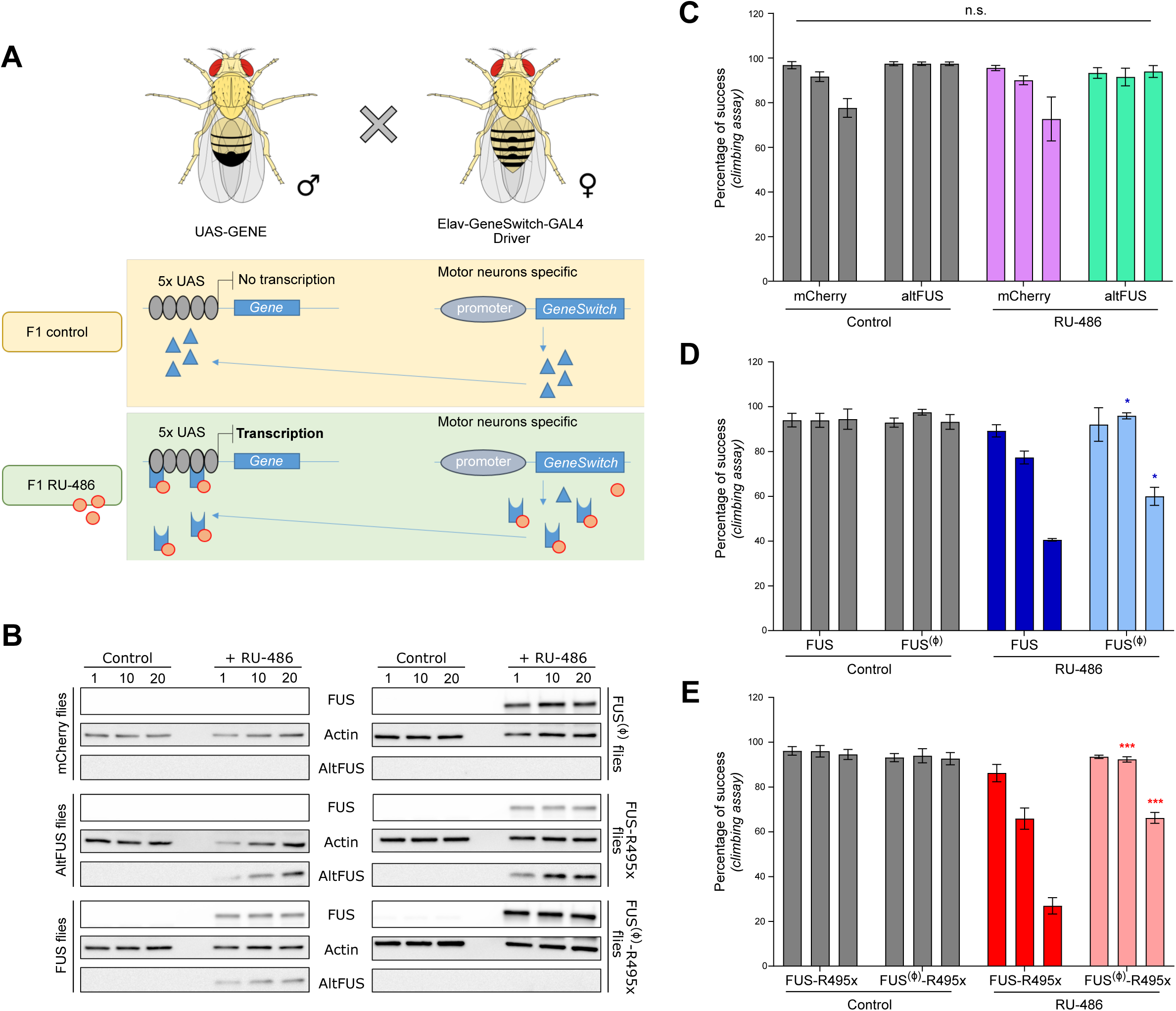
AltFUS expression is necessary for the full *FUS*-linked ALS phenotype in *Drosophila* **A**, Cross-breeding strategy for *Drosophila* generation using the Elav-GeneSwitch-GAL driver as an inducible expression system specific to the motor neurons. **B**, FUS and altFUS expression in mCherry (control), altFUS, FUS, FUS^(Ø)^, FUS-R495x or FUS^(Ø)^-R495x expressing *Drosophila* from the control F1 and the RU-486 treated F1 (see panel **A**) at 1, 10 or 20 days post-induction (representative image from n=2). **C-E**, Locomotion assay using the percentage of climbing success in control and RU-486-treated transgenic *Drosophila* expressing mCherry or altFUS (**C**), the bicistronic FUS or the monocistronic FUS^(Ø)^ (**D**), and the bicistronic FUS-R495x or the monocistronic FUS^(Ø)^-R495x (**E**) at day 1, 10 and 20 post-induction. Statistical comparison were made between each population (n=4). Indicated significance are between the monocistronic and the bicistronic transgenic flies of the RU-486-treated population (n.s. = non-significant, * = p value < 0.05, *** = p value < 0.001).

The motor neuron degeneration linked to ALS provokes a progressive locomotion loss measurable with a well-described climbing assay^52^. The control populations did not show any significant locomotion loss at day 1, 10 nor 20 (**Fig 5C-E**). Similarly to the RU-486 treated control group did not show a significant effect, although a decrease in climbing ability could be observed as previously reported^53, 54^ (mCherry transgenic flies – **Fig 5C**). AltFUS flies did not show any significant locomotion loss through time (**Fig 5C**). This result is consistent with the *in cellulo* data showing altFUS alone is not sufficient to provoke pathological hallmarks. As previously shown with this model^8^, the bicistronic FUS flies, which express both FUS and altFUS proteins, displayed a significant locomotion loss (**Fig 5D**). Bicistronic ALS-linked FUS-R495x flies showed an even greater motor neuron degeneration through time compared to FUS (**Fig 5E**). Monocistronic FUS^(Ø)^ (**Fig 5D**) and FUS^(Ø)^-R495x (**Fig 5E**) flies, which do not express altFUS, displayed both a delay in the onset of motor neuron degeneration and a reduced drop in climbing success at 20 days post-induction (40% vs. 60% for FUS^(Ø)^, 30% vs. 70% for FUS^(Ø)^-R495x). These results in *Drosophila* confirm a role for altFUS in *FUS*-related neurodegeneration *in vivo*, are consistent with our *in cellulo* observations and highlight the toxic cooperation between FUS and altFUS.

### ALS-ASSOCIATED MUTATIONS, SYNONYMOUS FOR FUS, ALTER ALTFUS AND LEAD TO TDP-43 CYTOPLASMIC AGGREGATES

As of today, over 50 mutations in the *FUS* gene have been associated with ALS^1^. However, most of these locate at the carboxyl end of the protein and as such have no effect on altFUS (**Fig 1A**).

We wondered whether mutations altering altFUS might have been overlooked as non-consequential in the FUS reading frame. We retrieved FUS synonymous mutations found in ALS patients, with an allelic frequency below 0.01 %, from previous studies and the ALS Variant Server (http://als.umassmed.edu/ - **Appendix Table 6**). The retrieved mutations clustered on the altFUS locus (**Fig 6A**), with 60 % of FUS synonymous mutations found in sALS patients and 50 % of FUS synonymous mutations found in fALS patients, which is significantly higher than expected by chance (34 %) (**Appendix Table 6**). We selected 4 mutations for further analysis based on the residue conservation: altFUS-P31L, altFUS-A38V, altFUS-A46V and altFUS-R64P. We generated them in GFP-FUS constructs: GFP-FUS^(P31L-FLAG)^-S44=; GFP-FUS^(A38V-FLAG)^-G51=; GFP-FUS^(A46V-FLAG)^-G59= and GFP-FUS^(R64P-FLAG)^-S77=. All altFUS mutants still localized to the mitochondria (**Fig EV8A**). To investigate whether these mutations may provoke an ALS-like phenotype, we quantified the number of cells presenting TDP-43 aggregates. All 4 altFUS mutants displayed clear TDP-43 aggregates, and showed a 1.8 to 2.4 fold increase compared to wild-type altFUS (**Fig 6B-C, Fig EV8B**). This result indicates that altFUS mutations potentiate TDP-43 cytoplasmic aggregation, a pathological hallmark in 98 % of ALS cases. Hence, some *FUS* mutations, synonymous for the FUS protein, exert a deleterious effect through their missense consequence on the altFUS protein.

**Figure 6:**
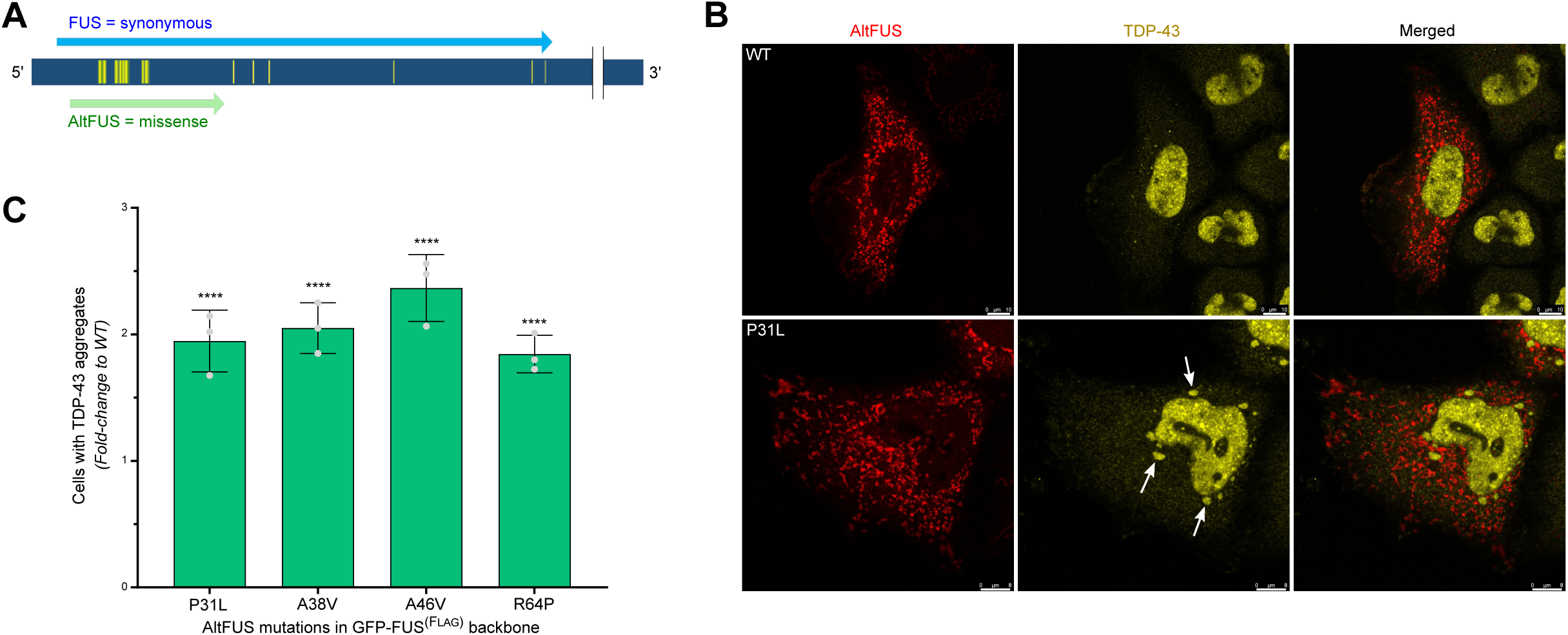
FUS mutations, synonymous for FUS but missense for altFUS, potentiate TDP-43 cytoplasmic aggregation **A**, Graphical representation of FUS synonymous mutations (yellow) found in ALS patients. The canonical FUS mRNA is represented in dark blue (*ENST00000254108* or *NM_004960*). The FUS protein coding sequence is indicated in light blue, and the altFUS protein coding sequence is indicated in green. **B**, Images by confocal microscopy of TDP-43 (yellow) and altFUS (FLAG tagged - red) in HeLa cells over-expressing GFP-FUS^(FLAG)^ or GFP-FUS^(P31L-FLAG)^-S44= (representative images from n=3). The white scale bar corresponds to 10 μm. **C**, Quantification of cells with TDP-43 aggregates in HeLa cells (see panel **B** and **Fig EV8B**). The data are represented as the fold-change compared to the GFP-FUS^(FLAG)^ expressing cells. Statistical significance is indicated above the bars (n=3, **** = p value < 0.0001).

### CONCLUSIONS: FUS AND ALTFUS, ENCODED BY THE SAME GENE, COOPERATE IN FUS-RELATED TOXICITY

Despite considerable advances in the field, current genome annotations still uphold arbitrary assumptions, such as the monocistronic nature of eukaryotic genes^16^. Here, we demonstrate *FUS* is a bicistronic gene. We discovered *FUS* CDS contains a second protein-coding sequence in a shifted frame overlapping its prion-like intrinsically disordered domain, regions known to host dual-coding events^22, 23^. This novel protein, named altFUS, is not an isoform but an entirely new sequence of 170 amino acids. AltFUS is endogenously expressed in human tissues and cultured cell lines, as demonstrated by ribosome profiling, mass spectrometry and with a custom antibody.

AltFUS is notably expressed in the motor cortex and iPSCs-derived motor neurons of healthy controls and ALS patients. Because altFUS is embedded within the *FUS* CDS, this discovery is of crucial importance to the field. Indeed, over-expression studies on *FUS* actually implicate two proteins: FUS and altFUS. Similarly FUS knockdown or knockout studies actually inhibit expression of both proteins^55^. Moreover, previous work has shown that gene editing techniques targeting a specific CDS do not necessarily result in knockout of the gene in case of dual-coding gene^56^. Our discovery thus suggests that *FUS* edited cells or models, notably targeting its last exons^57–59^, might only impair FUS protein expression but not altFUS, thus not resulting in a true *FUS* knockout. Our work provides a more accurate view of *FUS* coding potential to better understand its physiological function and models of FUS-related neurodegeneration.

Following the discovery of altFUS, we developed tools in order to differentiate the specific roles and phenotypes of FUS and altFUS. Our study demonstrates that altFUS is necessary for three toxic molecular hallmarks previously attributed to FUS: mitochondrial fragmentation and loss of mitochondrial membrane potential, inhibition of autophagy, and cytoplasmic aggregation of FUS. The inhibition of autophagy was observed when over-expressing altFUS alone, absent when over-expressing FUS alone (wild-type or ALS-associated R495x mutant) and reconstituted when co-expressing FUS and altFUS. This demonstrates that the inhibition of autophagy, previously described in *FUS*-ALS^10, 39^, has been incorrectly associated to the FUS protein. AltFUS inhibits autophagy. Moreover, altFUS is necessary but not sufficient for the mitochondrial membrane potential loss and cytoplasmic aggregation of FUS and TDP-43. Although TDP-43 aggregates are not commonly seen with FUS mutations^44^, it has already been reported^42, 44–49^ and this study lends support for altFUS conspiring with FUS and TDP-43 to lead to this molecular hallmark. Our data suggests the stoichiometry between FUS and altFUS may be important for the development of cytoplasmic aggregates. Both proteins are required to observe the phenotype, highlighting a functional alliance between FUS and altFUS. This pathological synergy was also observed in the *Drosophila* model.

The physiological function of altFUS is still unclear, although our work provides evidence for its role in mitochondrial dynamics and the cellular response to stress. One mechanism put forward in ALS is that the disease originates from a sub-optimal resolution of cellular stresses, which can come from environmental sources or mutated proteins^38, 60^. We have shown that altFUS, not FUS, inhibits autophagy, most likely via its interaction partners (**Appendix Table 5**). Since an inhibition of autophagy inhibits the dissociation of stress granules^38^, we suggest altFUS potentiates stress granule accumulation under stress conditions, and that FUS phase separation properties then lead to the formation of solid and toxic aggregates^61^. Further work is needed to fully understand the role of altFUS in physiological and pathological conditions, yet our study shows this novel protein plays a crucial role in *FUS*-linked gain-of-toxic dysfunctions in models of *FUS*-related neurodegeneration.

Recent studies have addressed the toxicity resulting from over-expression of wild-type FUS. Bogaert and colleagues used FUS domain truncation mutants to investigate wild-type FUS toxicity^51^. A FUS mutant lacking its N-terminal intrinsically disordered domain, thus lacking altFUS, displayed reduced toxicity. This study corroborates our findings that the absence of altFUS reduces the toxicity. Furthermore, Bogaert and colleagues concluded that FUS N-terminal synergizes with the C-terminal domain to mediate toxicity in *Drosophila*^51^. Here, we showed that altFUS synergizes with FUS to mediate ALS-like toxic features in cultured cells and toxicity in *Drosophila*. Despite different laboratories and different techniques, our data are in agreement with theirs. Yet, we point to an alternative (not necessarily mutually exclusive) explanation whereby altFUS, not the FUS N-terminal domain, synergizes for toxicity. Additionally, this discussion shows that not being aware of overlapping CDSs, especially in deletion studies, precludes alternative interpretations.

Current genome annotations guide the interpretation of data and the design of studies, however they also affect the way we screen for pathological mutations^16^. Most of the ALS-linked *FUS* mutations affect the carboxyl end of the protein, as shown with the example used in this study, the R495x mutation^62^. These mutations do not alter the altFUS protein, which is embedded at the beginning of the FUS CDS and span on exons 3 to 6. How could altFUS be important for the disease if most pathological mutations do not alter it? To answer this question, one needs to grasp how much genome annotations shape today’s research. For example, when screening for pathological mutations, those that are synonymous for FUS are discarded early in analyses as insignificant^63^. Yet, a synonymous mutation for FUS may not be for altFUS. Our work shows that FUS synonymous mutations found in patients cluster on altFUS genomic locus. We tested 4 of these mutations, synonymous for the FUS protein and missense for the altFUS protein, and we found that each of them potentiated TDP-43 cytoplasmic aggregation, a pathological hallmark of ALS^41^.

Overall, we have shown that *FUS* is a bicistronic gene and our results indicate that altFUS interferes with mitochondrial homeostasis and autophagy in cell culture and induces motor neuron toxicity in *Drosophila* models. It will be important to further characterize the function of altFUS and determine if it has a role in ALS and/or FTLD.

## METHODS

### FUS constructs

FUS and altFUS sequences were obtained from Bio Basic Gene Synthesis service. All FUS constructs were subcloned into pcDNA3.1-(Invitrogen) using Gibson assembly (New England Biolabs, E26115). FUS and altFUS wild-type sequences correspond to that of the Human FUS canonical transcript (*ENST00000254108* or *NM_004960*). FUS and altFUS proteins were tagged with V5 (GKPIPNPLLGLDST) and 2 FLAG (DYKDDDDKDYKDDDDK) respectively. FUS was tagged on the N-terminal, altFUS was tagged on the C-terminal. For immunofluorescence assays, N-terminal GFP-tagged FUS was also cloned into pcDNA3.1- by Gibson assembly. The necessary gBlocks were purchased from IDT. The monocistronic constructs FUS^(Ø)^ and FUS^(Ø)^-R495x were generated by mutating all altFUS methionines (ATG) to threonines (ACG). These mutations are synonymous in the FUS CDS (TAT > TAC, both coding for tyrosine). The altFUS mutated sequence was obtained from Bio Basic Gene Synthesis service, and then subcloned in FUS sequences in pcDNA3.1-using Gibson assembly. The bicistronic constructs are named as follows throughout the article: FUS, FUS-R495x, or FUS^(FLAG)^ and FUS^(FLAG)^-R495x when altFUS is FLAG-tagged in the +2 reading frame. The monocistronic constructs are named as follows throughout the article: FUS^(Ø)^ or FUS^(Ø)^-R495x to indicate altFUS absence.

### Cell culture, transfections, western blots and immunofluorescence

HEK293 and HeLa cells cultures tested negative for mycoplasma contamination (ATCC 30–1012K). Transfections, immunofluorescence, confocal analyses and western blots were carried out as previously described^64^. For FUS knock-down, 150 000 HEK293 cells in a 6-well plate were transfected with 25 nM FUS SMARTpool: siGENOME siRNA (Dharmacon, Canada, L-009497-00-0005) or ON-TARGET plus Nontargeting pool siRNAs (Dharmacon, D-001810-10-05) with DharmaFECT one transfection reagent (Dharmacon, T-2001–02) according to the manufacturer’s protocol. Cell media was changed every 24 hrs and cells were processed 72 hrs after transfection. For immunofluorescence, primary antibodies were diluted as follows: anti-Flag (Sigma, F1804) 1/ 1000, anti-TOMM20 (Abcam, ab186734) 1/500, anti-V5 (Cell Signalling Technologies, #13202) 1/1000, anti-TDP-43 (ProteinTech, 10782-2-AP) 1/500, and anti-TIAR (Cell Signalling Technologies, #8611) 1/1600. For western blots, primary antibodies were diluted as follows: anti-Flag (Sigma, F1804) 1/8000, anti-V5 (Sigma, V8012) 1/8000, anti-actin (Sigma, A5441) 1/40000, anti-FUS (Abcam, ab84078) 1/500, anti-altFUS (Abcam, custom antibody) 1/3000, anti-LC3 (Cell Signalling Technologies, #2775) 1/1000, anti-Hsp70 (Thermo Fisher Scientific, MA3-028) 1/1000, anti-Tubulin (Thermo Fisher Scientific, a11126) 1/2000 and anti-VDAC (Abcam, ab15895) 1/10000. The altFUS antibody was generated by injection two rabbits, each with 2 unique altFUS peptide. The purified antibody from rabbit 2 was used in this study at a 1/2000 dilution. Mitochondrial morphology was evaluated using the microP tool^65^. A minimum of 100 cells per replicate were counted across 3 independent experiments (n = 3, i.e. minimum 300 cells for each experimental condition). Colocalization analyses were performed using the JACoP plugin (Just Another Colocalization Plugin) implemented in Image J software, as previously described^13^. When specified, images obtained by confocal microscopy on the Leica TCS SP8 STED 3X were deconvolved using the Huygens software (Scientific Volume Imaging B.V., Hilversum, Netherlands). The software uses a signal reassignment algorithm for deconvolution, identical deconvolution parameters were applied to all images. The default parameters were used, including the Classic Maximum Likelihood Estimation (CMLE) algorithm, signal to noise ratio, and background estimation radius. The maximum iteration number was set at 30. Human tissue lysates for altFUS endogenous expression were purchased from Zyagen Laboratories (San Diego, California, USA).

### RIBO-seq data and conservation analyses

Global aggregate reads for initiating ribosomes and elongating ribosomes footprints across all available studies were downloaded from the Gwips portal (https://gwips.ucc.ie/), for *Homo sapiens* and for *Mus musculus*. For altFUS protein conservation analysis, all FUS mRNAs with at least EST evidence were retrieved across all available species from NCBI RefSeq. We performed an *in silico* 3-frame translation and retrieved the best matching protein sequence per species that displayed a minimum of 20 % sequence identity with the Human altFUS sequence over 25 % of Human altFUS length. AltFUS homologous sequences were found in 83 species, and we manually added that of *Drosophila melanogaster* that displayed a 37.5 % sequence identity over 19 % of the Human altFUS length. All retrieved altFUS sequences were then aligned using Clustalω with default parameters.

### Peptide-centric analysis of proteomics datasets

The stand alone PepQuery tool (v1.0) was downloaded from the PepQuery website (http://www.pepquery.org/). The tool was run on the following datasets from the TCGA consortium: colon cancer (COCA) proteome, ovarian cancer (OVCA) proteome, phosphoproteome and glycoproteome, and the breast cancer (BRCA) proteome and phosphoproteome. The reference database was set to the Ensembl datababase (hg38_Ensembl_20190910). The following parameters were set for all runs unless specified: carbamidomethylation of cysteine as fixed modification (as well as iTRAQ 4-plex of K, iTRAQ 4-plex of peptide N-term for BRCA and OVCA); oxidation of methionine as variable modification (as well as iTRAQ 4-plex of Y for BRCA and OVCA); a maximum of 3 modifications per peptides; trypsin digestion with maximum of 1 miscleavage; precursor tolerance of 10 ppm (20 ppm for COCA); fragment mass tolerance of 0.05 Da (0.6 Da for COCA); the hyperscore was used as a scoring metric, and 10 000 randoms. For phosphoproteomes, phosphorylation of Y, T and S were added as variable modifications. For the glycoproteome, deamidation of Q and N were added as variable modifications. PepQuery was run in the protein mode, with altFUS (IP_243680) whole sequence as input. Spectra were visualised and drawn using an in-house python script.

### Human induced pluripotent stem cell differentiation into motor neurons

Directed differentiation to human iPSC-motor neurons was performed as previously reported^27^. Briefly, iPSCs were maintained on Geltrex (Life Technologies) with Essential 8 Medium media (Life Technologies), and passaged using EDTA (Life Technologies, 0.5mM). All cell cultures were maintained at 37°C and 5% carbon dioxide. For motor neuron differentiation, iPSCs were differentiated to neuroepithelium by plating to 100% confluency in chemically defined medium consisting of DMEM/F12 Glutamax, Neurobasal, L-Glutamine, N2 supplement, non-essential amino acids, B27 supplement, β-mercaptoethanol (Life Technologies) and insulin (Sigma). Treatment with the following small molecules from day 0-7: 1µM Dorsomorphin (Millipore), 2µM SB431542 (Tocris Bioscience), and 3.3µM CHIR99021 (Miltenyi Biotec). At day 8, cells patterned for 7 days with 0.5µM retinoic acid and 1µM Purmorphamine. At day 14 spinal cord motor neuron precursors were treated with 0.1µM Purmorphamine for a further 4 days before being terminally differentiated for >10 days in 0.1 µM Compound E (Enzo Life Sciences) to promote cell cycle exit. Throughout the neural conversion and patterning phase (D0-18) the neuroepithelial layer was enzymatically dissociated twice (at D4-5 and D10-12) using dispase (GIBCO, 1 mg ml-1).

### Preparation of tissue lysates of the motor cortex of ALS patients

Approximately 100mg of motor cortex from 4 sporadic ALS and 4 C9orf72-ALS cases was lysed in 10x RIPA (50mM Tris HCl pH7.8, 150mM NaCl, 0.5% sodium deoxycholate, 1% NP40; supplemented with protease inhibitors and EDTA) volume using TissueLyzer equipment (Qiagen). Lysates were incubated on ice 20 minutes followed by centrifugation at 20,000xg for 20 minutes at 4°C. Supernatant was taken as ‘RIPA fraction’ and pellets resuspended in RIPA and SDS (final concentration of 2%). 3 sporadic ALS and 3 C9orf72-ALS samples were subsequently used as they were sufficiently concentrated to load 100 ug of proteins onto SDS-page gels.

### Mitochondrial extracts and cellular fractionation

Mitochondrial extracts were prepared as previously described^56^. Briefly, HEK293 cells grown up to 80 % confluence, were rinsed twice with PBS and gathered using a cell scraper. Cells were pelleted by centrifugation at 500 x g for 10 mins at 4 °C. Supernatant was discarded and cells suspended in mitochondrial buffer (210 mM mannitol, 70 mM sucrose, 1 mM EDTA, 10 mM HEPES-NaOH, pH 7.5, 0.5 mM PMSF and EDTA-free protease inhibitor (Thermo Fisher Scientific)). Cells were disrupted by 15 consecutive passages through a 25G1 0.5 x 25 needle syringe on ice, followed by a 3 min centrifugation at 2 000 x g at 4 °C. Supernatant was collected and the pellet suspended in mitochondrial buffer. The cell disruption was repeated four times and all retrieved supernatants containing mitochondria were again passed through syringe needle in mitochondrial buffer and cleared by centrifugation for 3 mins at 2000 × g at 4 °C. Supernatants were pooled and centrifuged for 10 mins at 13 000 x g at 4 °C to pellet mitochondria. The pellet was suspended in 200 μL of mitochondrial buffer until further processing. Cellular fractionation was performed using the Cell Fractionation Kit (#9038S, Cell Signaling Technology). Briefly, HEK293 cells were grown up to 80 % confluence, washed twice with PBS and gathered using a cell scraper. Cells were spun at 350 x g for 5 mins at 4 °C and 2.5 x 10^6^ cells were suspended in 500 μL of ice-cold PBS. An aliquot of 100 μL was spun at 350 x g for 5 mins at 4 °C and resuspended in SDS buffer (4 % SDS, Tris-HCl 100 mM pH 7.6) and kept as WCL (Whole Cell Lysate). The rest of the collected cells (remaining 400 μL) were spun at 500 x g for 5 mins at 4 °C. Supernatant was discarded and pellet resuspended in 500 μL of CIB (Cytoplasmic Isolation Buffer) from the kit, vortexed for 5 secs and incubated on ice for 5 mins. After centrifugation at 500 x g for 5 mins at 4 °C, the supernatant was collected as the cytosplasmic fraction. The pellet was resuspended in 500 μL of MIB buffer (Membrane Isolation Buffer) from the kit, vortexed for 15 secs and incubated on ice for 5 mins. After centrifugation at 8 000 x g for 5 mins at 4 °C, the supernatant was collected as the membrane and organelles fraction. To each 100 μL of fraction was added 60 μL of loading buffer 1 x (from ColdSpring Laemmli sample buffer: 50 mM Tris pH 6.8, 2 % SDS, 10 % Glycerol, 5 % β-mercaptoethanol) before processing for western blot.

### Mitochondrial membrane potential measurements

Mitochondrial membrane potential was measured by flow cytometry in HEK293 cells using TMRE (Tetramethylrhodamine, Ethyl ester, Abcam, ab113852). FCCP was used as a positive control to validate each independent experiment. Cells were grown up to 80 % confluence and washed twice with PBS. The cells were then incubated for 5 mins at 37 °C, 5 % CO_2_ with PBS/A (0.2 % BSA in PBS) solution (experimental) or 3 μM FCCP in PBS/A solution (positive control). Then, 100 nM of TMRE was added and cells were incubated 15 mins at 37 °C, 5 % CO_2_. After incubation, cells were trypsinized and centrifuged at 800 x g for 5 mins at 4 °C and resuspended in 500 μL of PBS and kept on ice. Cells were immediately analysed by flow cytometry. A gate for living cells was set, as well as a second gate to filter out cell doublets. TMRE fluorescence (PE-A) was recorded over a minimum of 50 000 gated cells for each experimental condition. The mean TMRE fluorescence intensity was measured over 3 independent experiments for each experimental condition.

### Stimulated Emission Depletion (STED) microscopy

Samples were prepared as described above for confocal microscopy. A Leica TCS SP8 STED 3X was used with a 100x objective lens and immersion oil for dual-color STED images. Images were obtained by sequential scanning of a given area. The combination of Alexa Fluor 488 (Thermo Fisher Scientific, A-11017) and Alexa Fluor 568 (Thermo Fisher Scientific, A-21069) dyes was chosen for STED imaging. Alexa Fluor 488 dye was excited with a white light laser (WLL) at 488 nm and was depleted using the 660 nm STED laser. Alexa Fluor 568 dye was excited with a WLL at 561 nm and was depleted using the 660 nm STED laser. The STED laser (660 nm) was applied at 80 % of maximum power.

### Fast Protein Liquid Chromatography (FPLC) and affinity-purification mass spectrometry (AP-MS)

Mitochondrial extracts of HEK293 cells were centrifuged at 13 000 x g for 10 mins at 4 °C to remove the supernatant and were resuspended in FPLC buffer (50 mM Tris-HCl, 1 mM EDTA, 150 mM NaCl, 1 % Triton X-100, pH 7.5, filtered with 0.2 μm filters) at 2 mg/mL for a total of 4 mg of mitochondrial proteins. Samples were incubated on ice for 15 mins and then centrifuged at 10 000 x g for 5 mins at 4 °C and the supernatant was loaded in the injection syringe without disrupting the pellet. The FPLC was performed on a HiLoad 16/60 Superdex 200 pg column (GE Healthcare, Chicago, USA) at 4 °C. The column was pre-equilibrated with the FPLC buffer for up to 0.2 CV (column volume) and the sample was applied at a flow rate of 0.5 mL/min with a pressure alarm set at 0.5 MPa. The elution was performed over 72 fractions of 1.5 mL for a maximum of 1.1 CV. For altFUS probing by western blot, proteins were precipitated from 150 μL of each 4 fractions in technical duplicates. First, 600 μL of methanol was added to each tube and mixed gently, before adding 150 μL of chloroform. Tubes were gently inverted 10 times before adding 450 μL of milliQ H_2_O and vortexing briefly. After centrifugation at 12 000 x g for 3 mins, the upper phase was discarded, and 400 μL of methanol was added. Tubes are centrifuged at 16 000 x g for 4 mins and the pellet was resuspended in loading buffer. For interactome analysis by mass spectrometry, fractions of interest (8 to 14) were pooled together and incubated at 4 °C overnight with magnetic FLAG beads (Sigma, M8823) pre-conditioned with FPLC buffer. The beads were then washed 3 times with 5 mL of FPLC buffer, and 5 times with 5 mL of 20 mM NH_4_HCO_3_ (ABC). Proteins were eluted and reduced from the beads using 10 mM DTT (15mins at 55 °C), and then treated with 20 mM IAA (1 hour at room temperature in the dark). Proteins were digested overnight by adding 1 μg of trypsin (Promega, Madison, Wisconsin) in 100 μL ABC at 37 °C overnight. Digestion was quenched using 1 % formic acid and supernatant was collected. Beads were washed once with acetonitrile/water/formic acid (1/1/0.01 v/v) and pooled with supernatant. Peptides were dried with a speedvac, desalted using a C18 Zip-Tip (Millipore Sigma, Etobicoke, Ontario, Canada) and resuspended into 30 μl of 1% formic acid in water prior to MS analysis.

### Mass-spectrometry analysis

Peptides were separated in a PepMap C18 nano column (75 μm × 50 cm, Thermo Fisher Scientific). The setup used a 0–35% gradient (0–215 min) of 90% acetonitrile, 0.1% formic acid at a flow rate of 200 nL/min followed by acetonitrile wash and column re-equilibration for a total gradient duration of 4 h with a RSLC Ultimate 3000 (Thermo Fisher Scientific, Dionex). Peptides were sprayed using an EASYSpray source (Thermo Fisher Scientific) at 2 kV coupled to a quadrupole-Orbitrap (QExactive, Thermo Fisher Scientific) mass spectrometer. Full-MS spectra within a m/z 350–1600 mass range at 70,000 resolution were acquired with an automatic gain control (AGC) target of 1e6 and a maximum accumulation time (maximum IT) of 20 ms. Fragmentation (MS/MS) of the top ten ions detected in the Full-MS scan at 17,500 resolution, AGC target of 5e5, a maximum IT of 60 ms with a fixed first mass of 50 within a 3 m/z isolation window at a normalized collision energy (NCE) of 25. Dynamic exclusion was set to 40 s. Mass spectrometry RAW files were searched with Andromeda search engine implemented in MaxQuant 1.5.5.1. The digestion mode was set at Trypsin/P with a maximum of two missed cleavages per peptides. Oxidation of methionine and acetylation of N-terminal were set as variable modifications, and carbamidomethylation of cysteine was set as fixed modification. Precursor and fragment tolerances were set at 4.5 and 20 ppm respectively. Files were searched using a target-decoy approach against UniprotKB (*Homo sapiens* 03/2017 release) with the addition of altFUS sequence for a total of 92,949 entries. The false discovery rate (FDR) was set at 1% for peptide-spectrum-match, peptide and protein levels. Protein interactions were then scored using the SAINT algorithm, with Mock cells as control and the magnetic FLAG beads in HEK293 cells crapome^66^. Proteins with a SAINT score above 0.99 were considered, as well as those presenting a SAINT score above 0.88 with a minimum of two unique peptides.

### Biological processes and cellular compartment enrichment analysis

Proteins identified in altFUS interactome were screened for cellular compartment and biological processes enrichment using Gene Ontology (GO) enrichment. Proteins were queried against the whole Human Proteome for cellular compartment and against the Human mitochondrial proteome (MitoCarta 2.0) for biological processes. The statistical analysis used a Fisher’s Exact test with a FDR set at 1 %.

### Autophagic flux measurements

The mCherry-GFP-LC3 was used to evaluate the autophagic vesicles within HeLa cells by confocal microscopy. Before fusion with the lysosome, the LC3 molecules on the autophagosome display a yellow fluorescence (combined mCherry and GFP fluorescence). After fusion, the GFP fluorescence is quenched by the lysosomal pH, and as such the LC3 molecules display a red signal (mCherry alone). This allows a visual representation of the autophagic flux in a given cell. Cells treated with 50 nM Bafilomycin for 4 hours were used as a positive control to validate each independent experiment. Observations were made across 2 technical duplicates for each biological condition, across 3 independent experiments (n=3). Alternatively, the autophagic flux was also evaluated by LC3 probing before and after bafilomycin treatment (50 nM for 4 hours). The quantification corresponds to the treated / untreated ratio of LC3-II abundance.

### Cytoplasmic aggregates measurements

Images of HeLa cells were taken by confocal microscopy and then processed using the Image J 3D Objects Counter plugin. FUS cytoplasmic aggregates were then quantified in number and size (μm^2^) for each cell. A total of 100 cells across two technical replicates were taken for each independent experiment (n=3, i.e. a minimum of 300 cells per biological conditions).

### Transgenic *Drosophila* and climbing assay

The bicistronic constructs, FUS and FUS-R495x, and the monocistronic constructs, altFUS, FUS^(Ø)^ and FUS^(Ø)^-R495x, were subcloned in the pUASTattB expression vector for site specific insertion into attP2 on chromosome 3. Transgenic flies were generated by Best Gene (Best Gene Inc., California, USA). The Elav-GeneSwitch-GAL4 driver (stock number: 43642, genotype: y[1] w[*]; P{w[+mC]=elav-Switch.O}GSG301) and the UAS-mCherry flies (stock number: 35787, genotype: y[1] sc[*] v[1]; P{y[+t7.7] v[+t1.8]=UAS-mCherry.VALIUM10}attP2) was purchased from Bloomington (Bloomington Drosophila Stock Center, Indiana, USA). All stocks were in a *w^1118^* background and were cultured on standard medium at 25°C or room temperature. Transgenic flies were crossed with the Elav-GeneSwitch-GAL4 driver strain. The F1 was equally divided in two groups with equal proportion of males and females: one group will feed on standard food supplemented with ethanol (0.2 % - control flies), the other on standard food supplemented with RU-486 at 10 μM diluted in ethanol (induced flies). The climbing assay was performed as previously described^52^. Briefly, flies were transferred into an empty vial and tapped to the bottom. After 18 s, the number of flies at the top of the tube were considered successful. The assay was done at day 1, 10 and 20 post-induction, across 4 independent F1. Five flies were taken at day 1, 10 and 20 post-induction to validate expression of the proteins of interest.

### Statistical analyses and representation

Unless otherwise stated, the statistical analysis carried was a two-way ANOVA with Tukey’s multiple comparison correction. The box plots represent the mean with the 5 to 95 % percentile. The bar graphs represent the mean, and error bars correspond to the standard deviation. When using parametric tests, normality of data distribution was verified beforehand using the Shapiro-Wilk test.

### DATA AVAILABILITY STATEMENT

The OpenProt database is available at www.openprot.org. The GTEx portal is available via www.gtexportal.org. The Gwips portal is available at www.gwips.ucc.ie. The proteomics data are available on the PRIDE repository with the accession PXD------. Any other relevant data are available from the corresponding authors upon reasonable request.

## ACKNOWLEDGEMENTS

This research was supported by CIHR grants MOP-137056 and MOP-136962, by an ALS Canada Project Grant, and by a Canada Research Chair in Functional Proteomics and Discovery of Novel Proteins to X.R. X.R and S.J are members of the Fonds de Recherche du Québec Santé (FRQS)-supported Centre de Recherche du Centre Hospitalier Universitaire de Sherbrooke. P.M. is supported by the ALS Double Play Christopher Chiu Postdoctoral Fellowship. We thank J. Ule, M-J. Boucher and D. Hunting for helpful discussions. The autopsy program is supported by the James Hunter and Family ALS Initiative. The Genotype-Tissue Expression (GTEx) Project was supported by the Common Fund of the Office of the Director of the National Institutes of Health, and by NCI, NHGRI, NHLBI, NIDA, NIMH, and NINDS. The data used for the analyses described in this manuscript were obtained from the GTEx Portal on April 2019.

## CONTRIBUTIONS

MA.B and X.R designed and wrote the study. MA.B and JF.J did the experiments, and MA.B did the analyses and figures. S.N and S.J assisted with the Drosophila experiments. P.M, L.Z and J.R provided the motor cortex tissue lysates. GE.T and R.P provided the iPSCs-derived motor neurons. All authors proofread the manuscript.

## LEGENDS

### Extended View Figures

**Figure EV1:**
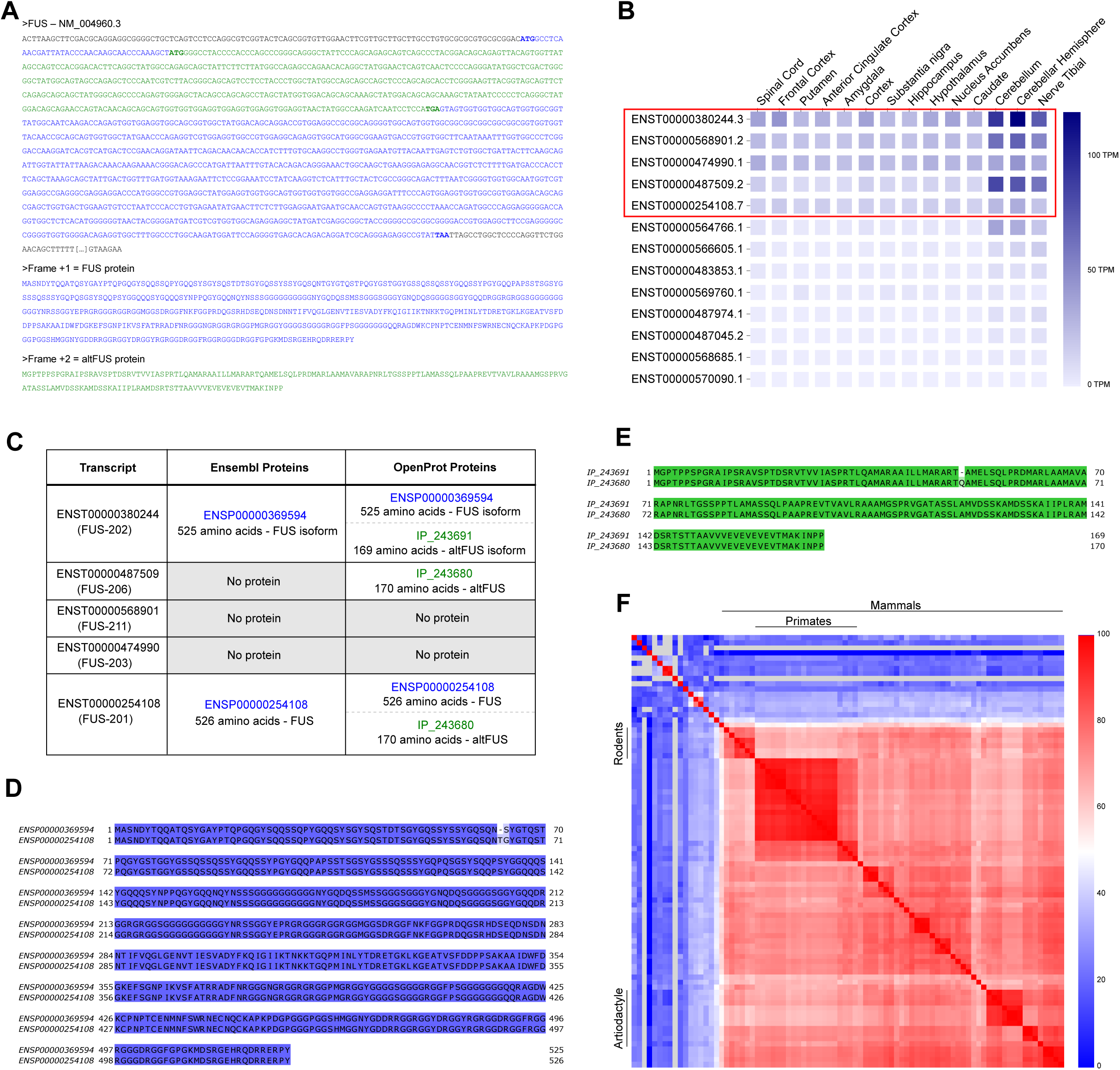
*FUS* gene protein diversity **A**, AltFUS is encoded within an alternative open reading frame (ORF) overlapping the FUS canonical CDS. When read in the +1 frame, the *FUS* mRNA (here *ENST00000254108*) codes for a 526 amino acid protein (highlighted in blue), named FUS. In the +2 frame, the *FUS* mRNA contains a second ORF (highlighted in green) that codes for a novel 170 amino acid protein, named altFUS. The two proteins are not isoforms. **B**, GTEx portal data on *FUS* mRNA expression in the brain and nerves are shown in blue colored scale (TPM = Transcripts Per Million). Transcripts are identified with Ensembl accessions (the number after the dot corresponds to the version used in the analysis) and are quantified across 14 tissues. The five transcripts framed in red share 85 % of all mRNA expression level. **C**, Table compiling the protein information relayed by Ensembl and OpenProt resources for the five transcripts highlighted in panel **B**. The FUS protein is highlighted in blue. The altFUS protein is highlighted in green. D-E, Alignment (Clustalω) of protein sequences for FUS (ENSP00000254108) and its isoform (ENSP00000369594) in panel **D** (blue) and altFUS (IP_243680) and its isoform (IP_243691) in panel **E** (green). The residues are coloured based on their degree of identity. **F**, Heatmap of altFUS protein sequences identity across 84 species (see **Appendix Table 2** and **Source Data 1**). Primates, Rodents and Mammals in general display strong protein conservation. The sequence identity is colored from blue (0 %) to red (100 %).

**Figure EV2:**
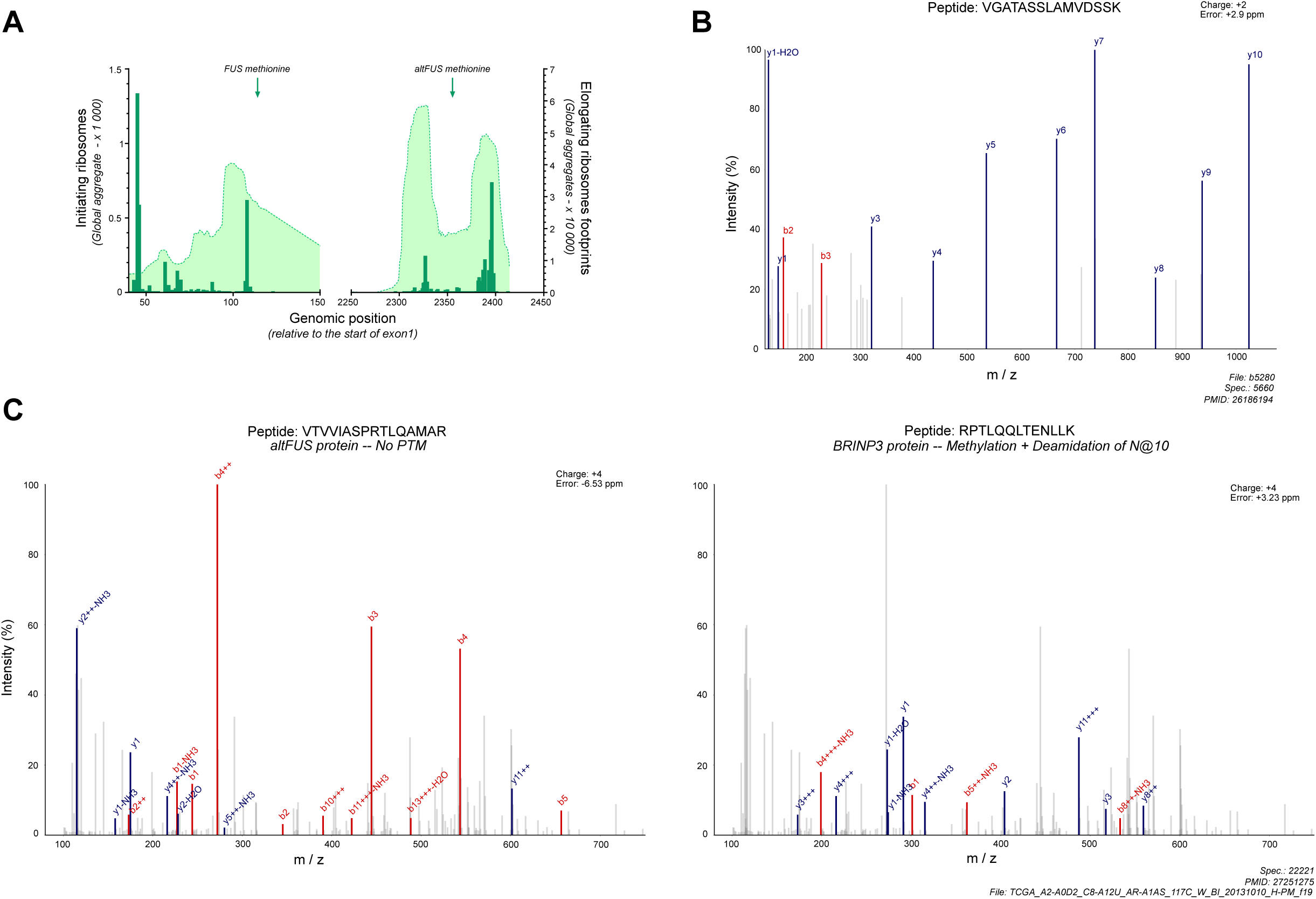
AltFUS is endogenously expressed **A**, RIBO-seq data over the mouse *FUS* gene from the Gwips portal (*Mus musculus*). Initiating ribosome reads are indicated by green bars, and elongating ribosomes footprints are indicated by the green curve. The graph capture the beginning of the *FUS* gene with FUS and altFUS methionines indicated by green arrows. The genomic positions are indicated relative to the start of exon 1. **B**, MS/MS spectra confidently mapped to a peptide unique to altFUS by OpenProt resource. Peaks are represented by their mass over charge ratios (m/z) and their intensity relative to the highest (relative intensity). The y ions are coloured in blue, the b ions in red and the unannotated peaks appear in grey. The charge state and m/z error of the matched peptide are indicated on the top right corner of the spectra. The original study, file and spectra number are indicated below the graph. **C**, Comparison of one MS/MS spectra confidently mapped to a peptide unique to altFUS (left spectra) and its best possible annotation with any known protein with any post-translational modification (PTM - right spectra). Peaks are represented by their mass over charge ratios (m/z) and their intensity relative to the highest (relative intensity). The y ions are coloured in blue, the b ions in red and the unannotated peaks appear in grey. The charge state and m/z error for each of the matched peptides are indicated on the top right corner of each panel. The original study, file and spectra number are indicated below the graph. On the top of each panel is indicated the matched spectra with its corresponding protein and PTMs.

**Figure EV3:**
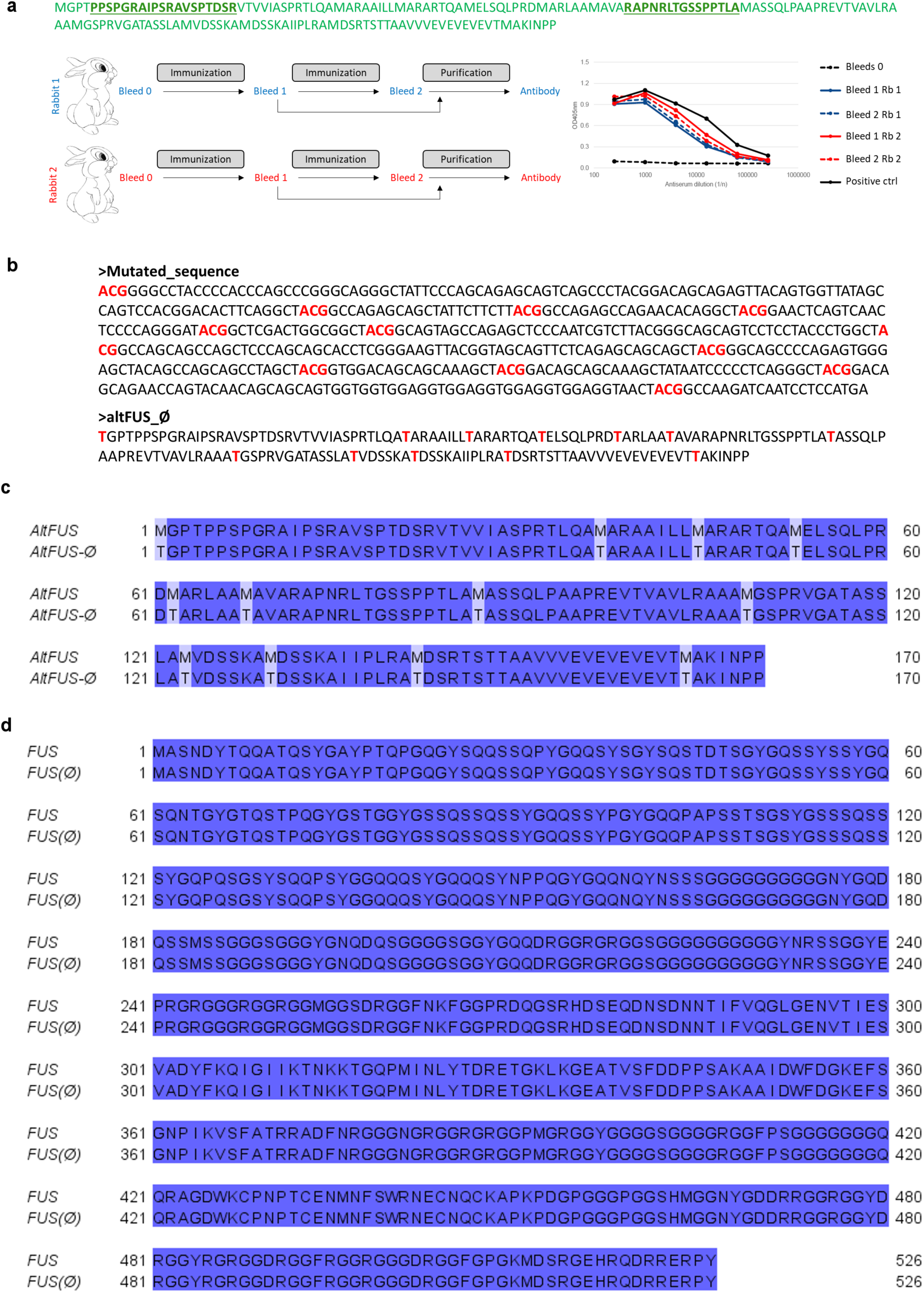
Tools to explore altFUS expression and roles **A**, AltFUS custom antibody strategy, with epitopes highlighted in bold on altFUS sequence (green), and ELISA test results on both bleeds from each rabbits. B, Nucleotidic mutations on *FUS* mRNA (*ENST00000254108*) are highlighted in red and mutate all altFUS methionines to threonines (altFUS_Ø). **C-D**, Protein alignment of altFUS (**C**) and FUS (**D**) proteins from FUS bicistronic construct and the monocistronic (FUS^(Ø)^) construct. The residues are coloured based on sequence identity (dark blue for identical residues).

**Figure EV4:**
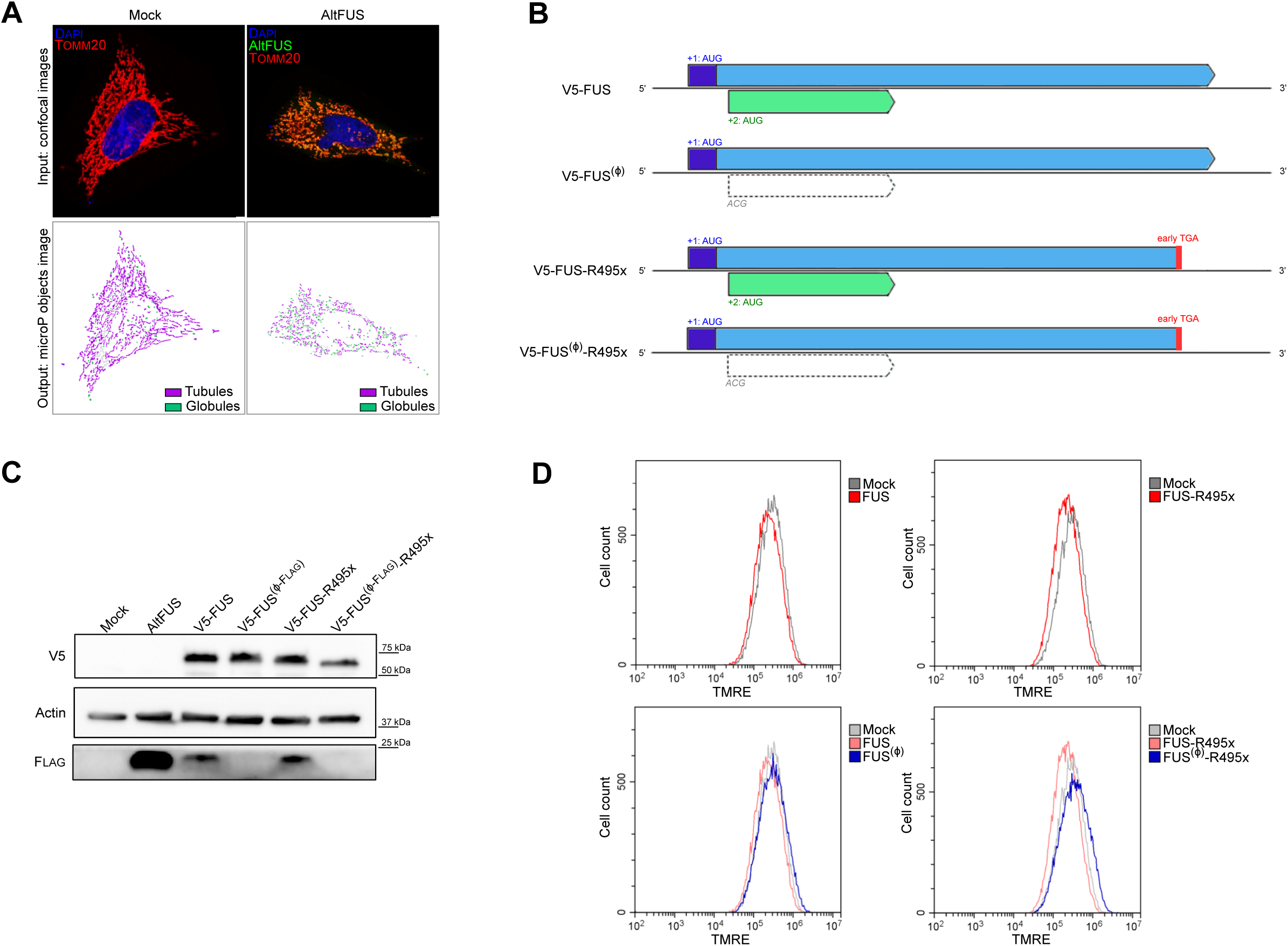
AltFUS is involved in mitochondrial functions **A**, MicroP processing of confocal images of mock and altFUS transfected HeLa cells. The nucleus is coloured in blue, altFUS in green and the mitochondrial network in red (TOMM20). Tubules are coloured in purple, globules are coloured in green. **B**, Graphical representation of bicistronic constructs FUS and FUS-R495x (early stop codon highlighted in red), and their monocistronic equivalent FUS^(Ø)^ and FUS^(Ø)^-R495x. AltFUS CDS is highlighted in green when present, FUS CDS is highlighted in blue. **C**, V5-FUS, V5-FUS-R495x and altFUS-FLAG expression from bicistronic or monocistronic constructs in HEK293 cells (representative image from n=3). **D**, Representative traces of TMRE fluorescence measured by flow cytometry in mock cells and cells overexpressing the bicistronic constructs FUS or FUS-R495x (in red) in the top panels, and the monocistronic constructs, FUS^(Ø)^ or FUS^(Ø)^-R495x (in blue) in the bottom panels (n=3, minimum of 50 000 live cells per independent replicates).

**Figure EV5:**
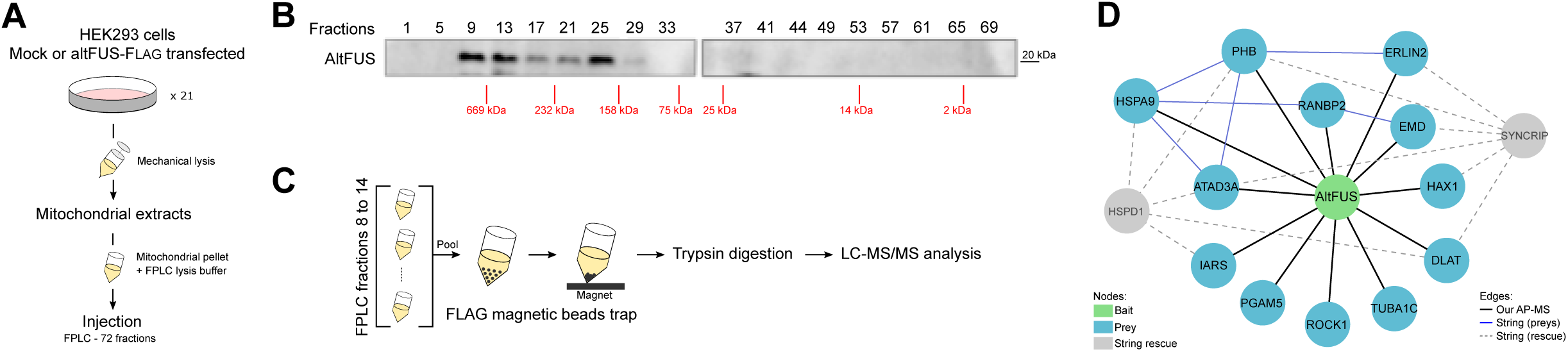
AltFUS mitochondrial interactome **A**, Workflow of mitochondrial extraction and size exclusion chromatography (FPLC) on mock or altFUS-FLAG transfected HEK293 cells. **B**, AltFUS expression in FPLC fractions on transfected HEK293 cells mitochondrial extracts (representative image from n=2). Every 4 fractions were loaded, the size scale through the fractions is indicated in red. **C**, Workflow of magnetic beads FLAG trap on FPLC fractions 8 to 14, from mock or altFUS-FLAG transfected HEK293 cells. **D**, Protein-protein interaction network of altFUS (bait, in green). Identified proteins (preys) are indicated in blue. The network was expanded using String database, with a maximum of 2 additional nodes (grey nodes and grey dotted edges). Interactions (edges) identified by our AP-MS are in black. Interactions between preys were added using the String database (blue edges).

**Figure EV6:**
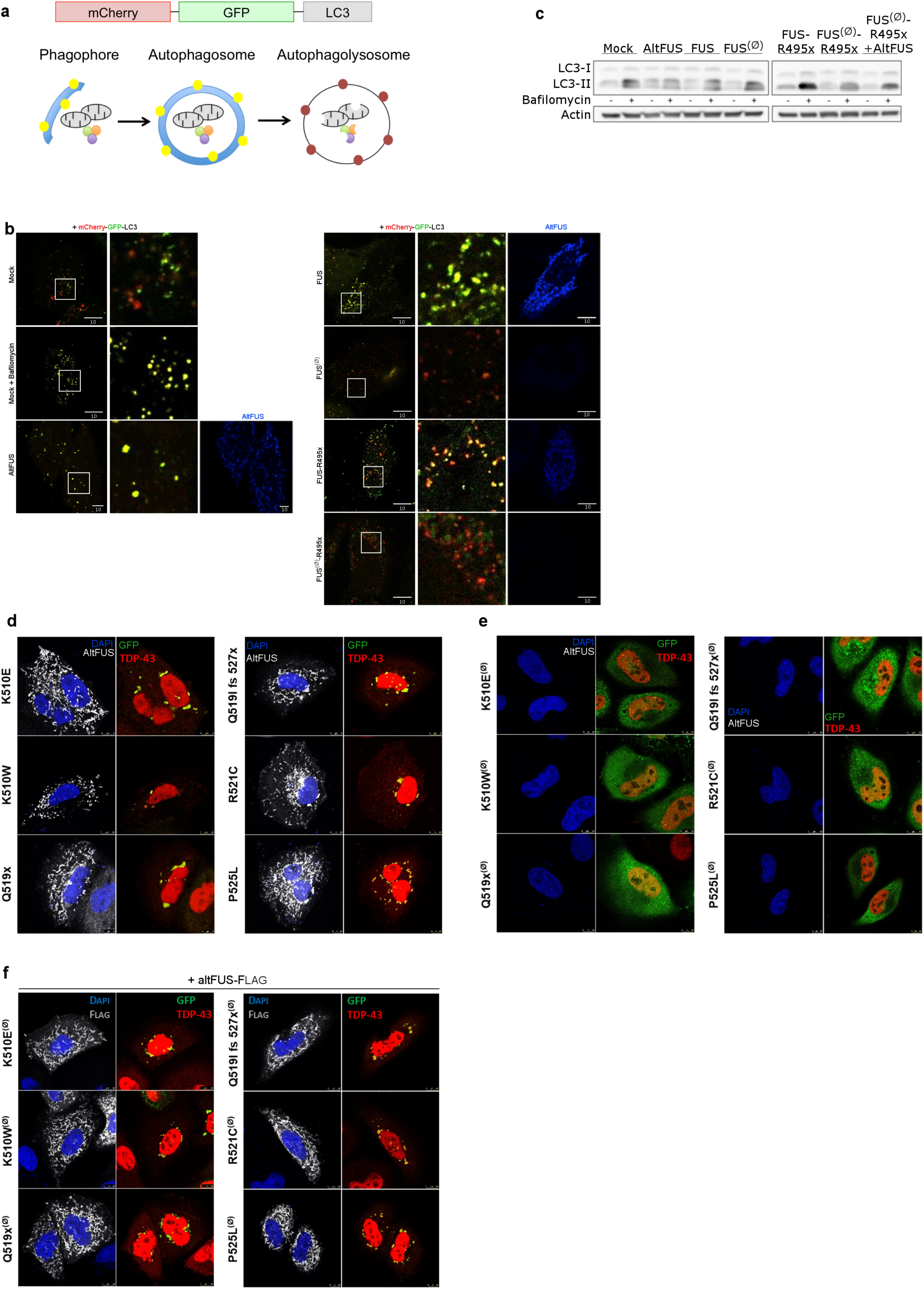
AltFUS inhibits the autophagy and potentiate FUS/TDP-43 cytoplasmic aggregates **A**, Graphical representation of the mCherry-GFP-LC3 construct to study the autophagic flux. **B**, Images by confocal microscopy of mCherry-GFP-LC3 and altFUS (FLAG, in blue) signal in HeLa cells across biological conditions: untreated mock, bafilomycin treated mock, altFUS, FUS, FUS^(Ø-FLAG)^, FUS-R495x and FUS^(Ø-FLAG)^-R495x (representative images of n=3). The white scale bar corresponds to 10 μm and the zoomed in region (right panel) is delimited as a white box (related to **Fig4A**). **C**, LC3-II expression from untreated (-) or bafilomycin treated (+) cells transfected with mock, altFUS, FUS, FUS^(Ø)^, FUS-R495x, FUS^(Ø)^-R495x, or co-transfected with FUS^(Ø)^-R495x and altFUS. **D-F**, Images by confocal microscopy of FLAG (white), GFP (green) and TDP-43 (red) signals in HeLa cells transfected with the bicistronic constructs (**D**), monocistronic constructs (**E**) or co-transfected with altFUS-FLAG and the monocistronic constructs (**F**) of 6 ALS-associated mutants: FUS-G156E, FUS-K510E, FUS-Q519x, FUS-Q519I-fs527x, FUS-R521C, and FUS-P525L (representative images from n=3). The white scale bar corresponds to 10 μm.

**Figure EV7:**
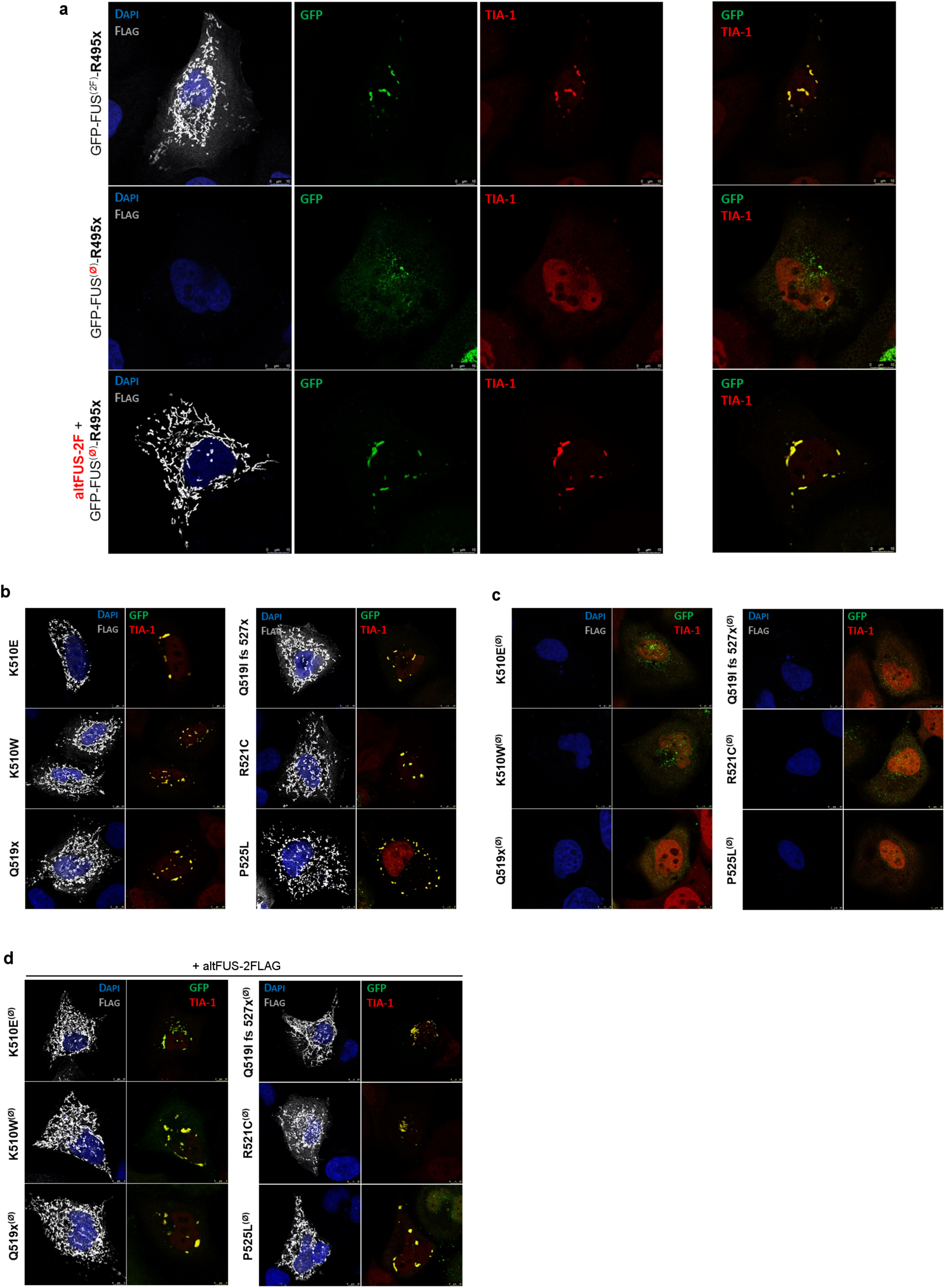
AltFUS potentiate FUS recruitment to stress granules **A**, Images by confocal microscopy of FLAG (white), GFP (green) and TIA-1 (red) signals in HeLa cells transfected with the bicistronic GFP-FUS^(FLAG)^-R495x or the monocistronic GFP-FUS^(Ø-FLAG)^-R495x constructs, or co-transfected with the monocistronic GFP-FUS^(Ø-FLAG)^-R495x and altFUS-FLAG constructs (representative images from n=3). The white scale bar corresponds to 10 μm. **B-D**, Images by confocal microscopy of FLAG (white), GFP (green) and TIA-1 (red) signals in HeLa cells transfected with the bicistronic constructs (**B**), monocistronic constructs (**C**) or co-transfected with altFUS-FLAG and the monocistronic constructs (**D**) of 6 ALS-associated mutants: FUS-G156E, FUS-K510E, FUS-Q519x, FUS-Q519I-fs527x, FUS-R521C, and FUS-P525L (representative images from n=3). The white scale bar corresponds to 10 μm.

**Figure EV8:**
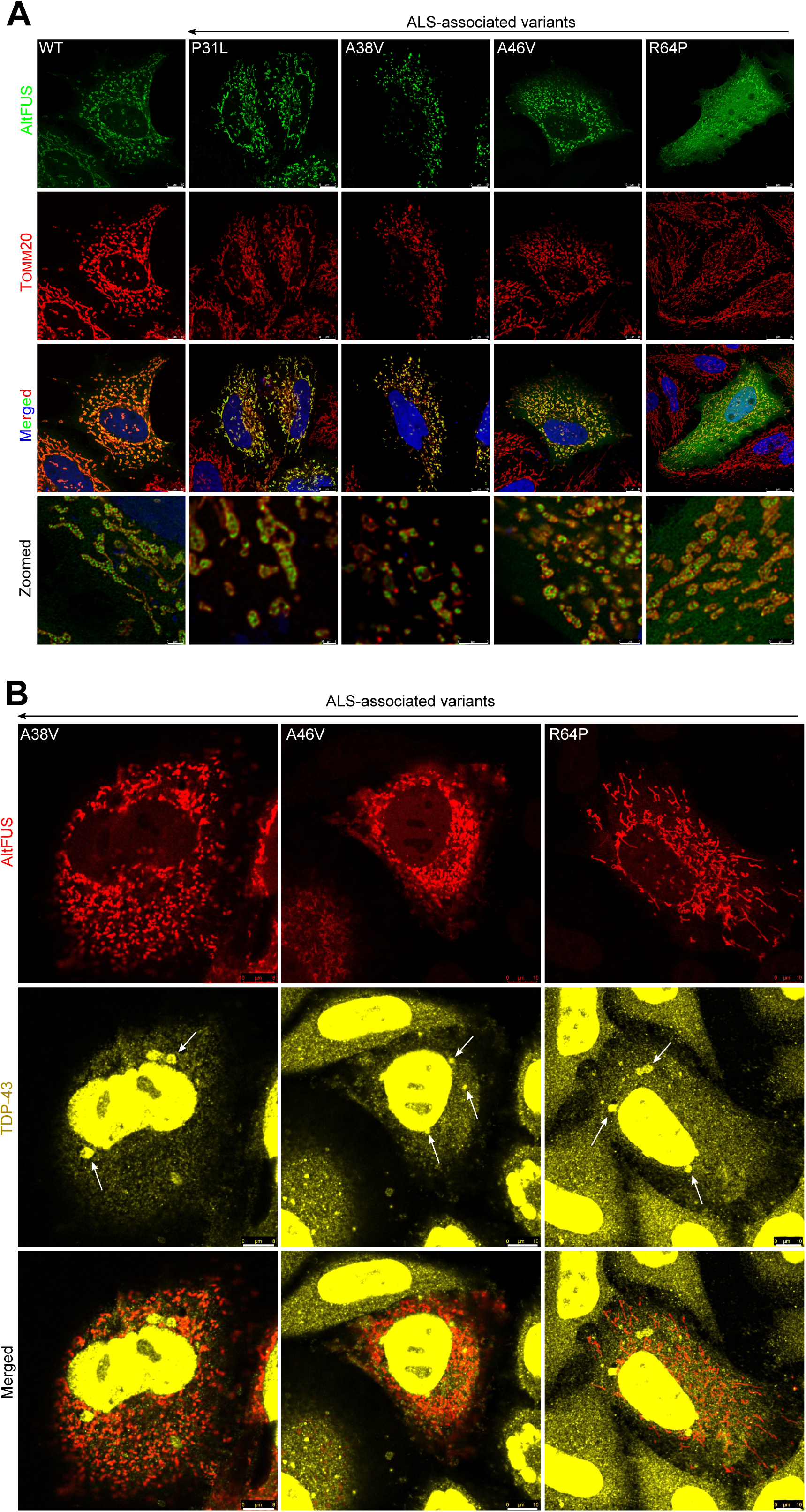
Mitochondrial altFUS mutants potentiates TDP-43 cytoplasmic aggregates **A**, Images by confocal microscopy of altFUS (FLAG tagged - green) and mitochondria (TOMM20 marker, red) in HeLa cells over-expressing altFUS-FLAG, altFUS-P31L-FLAG, altFUS-A38V-FLAG, altFUS-A46V-FLAG or altFUS-R64P-FLAG constructs (representative images from n=3). The white scale bar corresponds to 10 μm. Deconvolution over a maximum of 30 iterations on the green and red channels was performed for the zoomed in pictures. The white scale bar corresponds to 3 μm. **B**, Images by confocal microscopy of TDP-43 (yellow) and altFUS (FLAG tagged - red) in HeLa cells over-expressing GFP-FUS^(A38V-FLAG)^-G51=, GFP-FUS^(A46V-FLAG)^-G59= or GFP-FUS^(R64P-FLAG)^-S77= constructs (representative images from n=3). The white scale bar corresponds to 10 μm.

### Appendix Tables

**Appendix Table 1: *FUS* gene annotation on OpenProt OpenProt search results for FUS transcripts *ENST00000254108* and *NM_004960.3*.** The OpenProt version 1.3 was used. Each protein prediction is identified with a unique protein accession (starting with II_ for novel predicted isoform, and IP_ for predicted alternative proteins). All predicted alternative proteins are coloured in light blue. The protein localization is relative to the canonical FUS coding sequence (CDS). The furthered studied protein, named altFUS (IP_243680) is highlighted in dark blue. The workbook also contains the list of peptides identified in re-analysis of mass spectrometry experiments using the OpenProt database (version 1.3).

**Appendix Table 2: altFUS protein sequences across species AltFUS conservation across vertebrates.** The workbook contains the list of *FUS* transcripts retrieved from NCBI RefSeq database (transcript accession and nucleotidic sequences); the list of transcripts containing an altFUS sequence (transcript accession, species, and altFUS protein sequence); and the matrix of altFUS protein sequence identity between each species.

**Appendix Table 3: Peptide-Centric proteomic results for altFUS endogenous expression Peptide-spectrum matches confidently identified using PepQuery (v1.0) on the TCGA datasets.** The first sheet of the workbook contains the legends for the summary and results tables. The summary table lists the number of peptides and PSMs in each dataset. The details of results for each dataset queried is also listed.

**Appendix Table 4: Post-translational modification predictions on altFUS sequence Predictions for post-translational modifications (PTMs) on altFUS sequence.** The summary sheets contains the name of the modification predicted, the target residues, the tool used and the number of predicted sites with their confidence score. Results for the 2 more abundant PTMs predicted (phosphorylation and O-Glc-NAcylation) are detailed.

**Appendix Table 5: AltFUS interacting proteins Proteins identified by mass spectrometry following size exclusion chromatography (FPLC) and FLAG affinity purification on mitochondrial extracts from HEK293 cells over-expressing altFUS-FLAG.** The proteins are listed by their UniProtKB accession alongside their annotated function, the number of unique peptides supporting the detection and the number of peptide spectrum matches (PSMs) within each experimental condition. Confidence of the interaction was scored using the SAINT algorithm. The SAINT score is also reported. Proteins involved in autophagy related processes are indicated in red. In light greys are proteins removed from the analysis because lacking protein coverage (unique peptides < 2).

**Appendix Table 6: FUS synonymous mutations in ALS patients and their consequence on altFUS *FUS* mutations, synonymous for FUS but missense for altFUS.** The first sheet contains mutations identified from a manually curated literature review. The second sheet retrieves variants in sALS cases from the ALS Variant Server (http://als.umassmed.edu/). The third sheet retrieves variants in fALS cases from the ALS Variant Server.

### Source Data

**Source Data 1:**
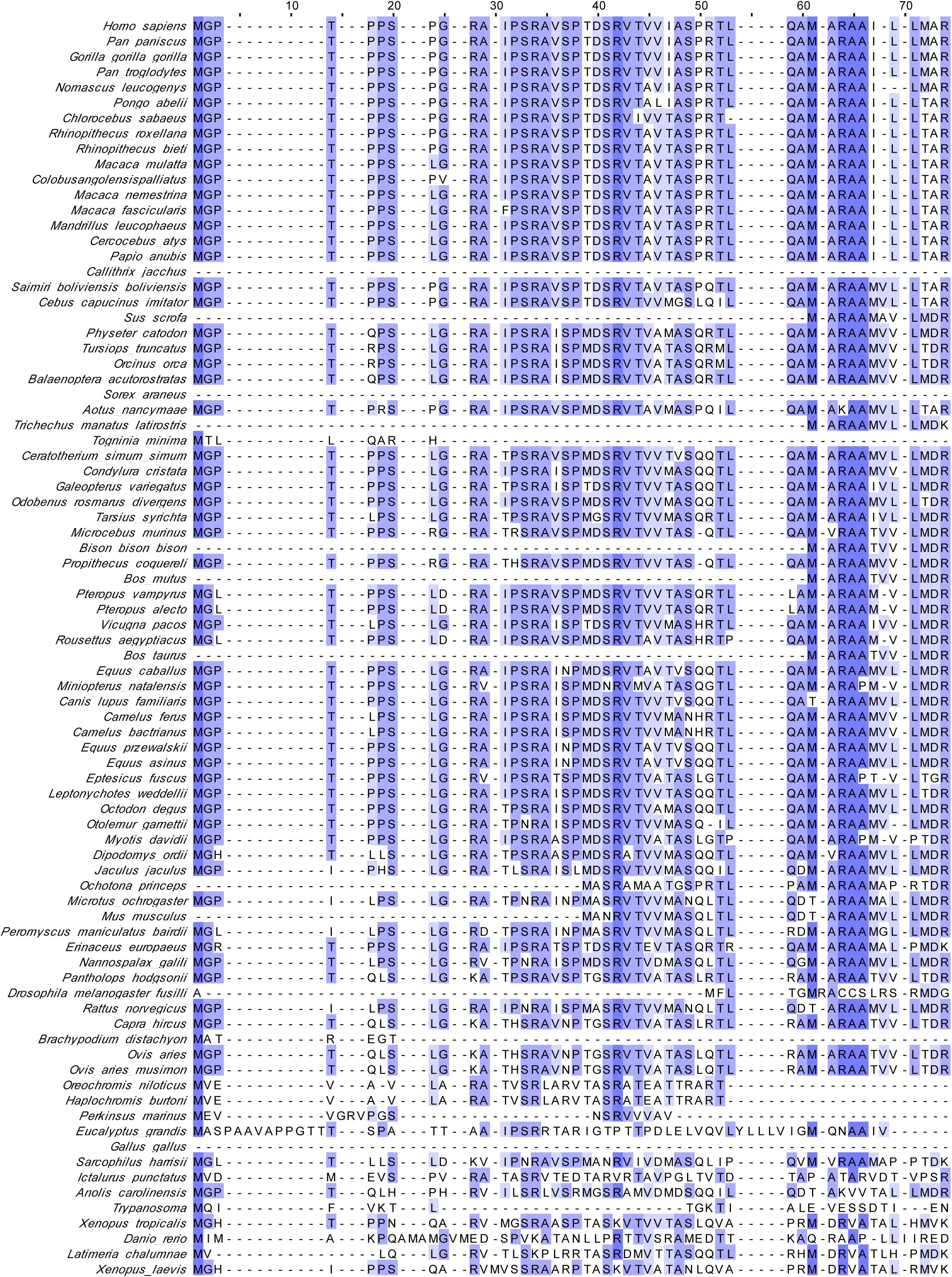
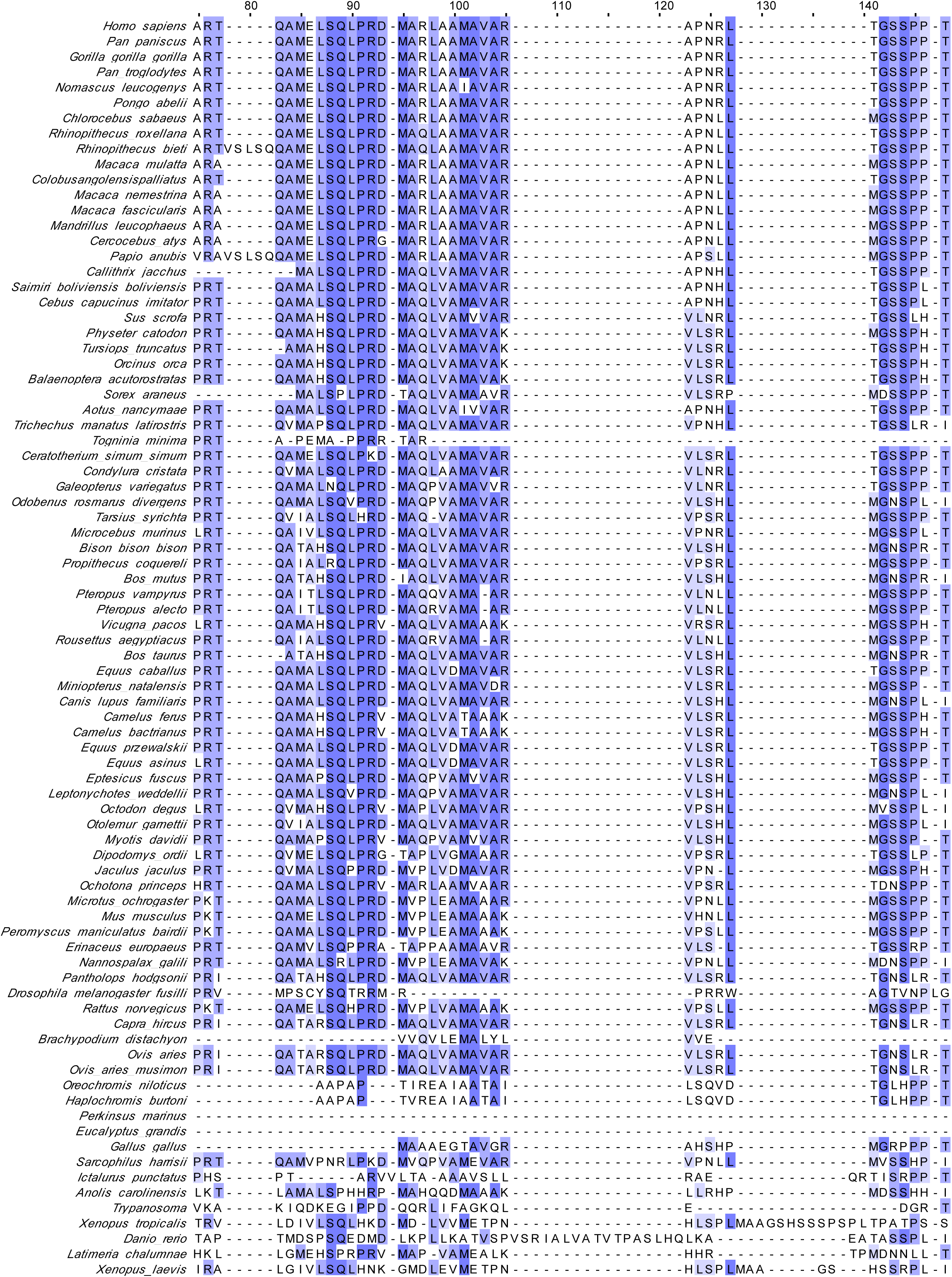
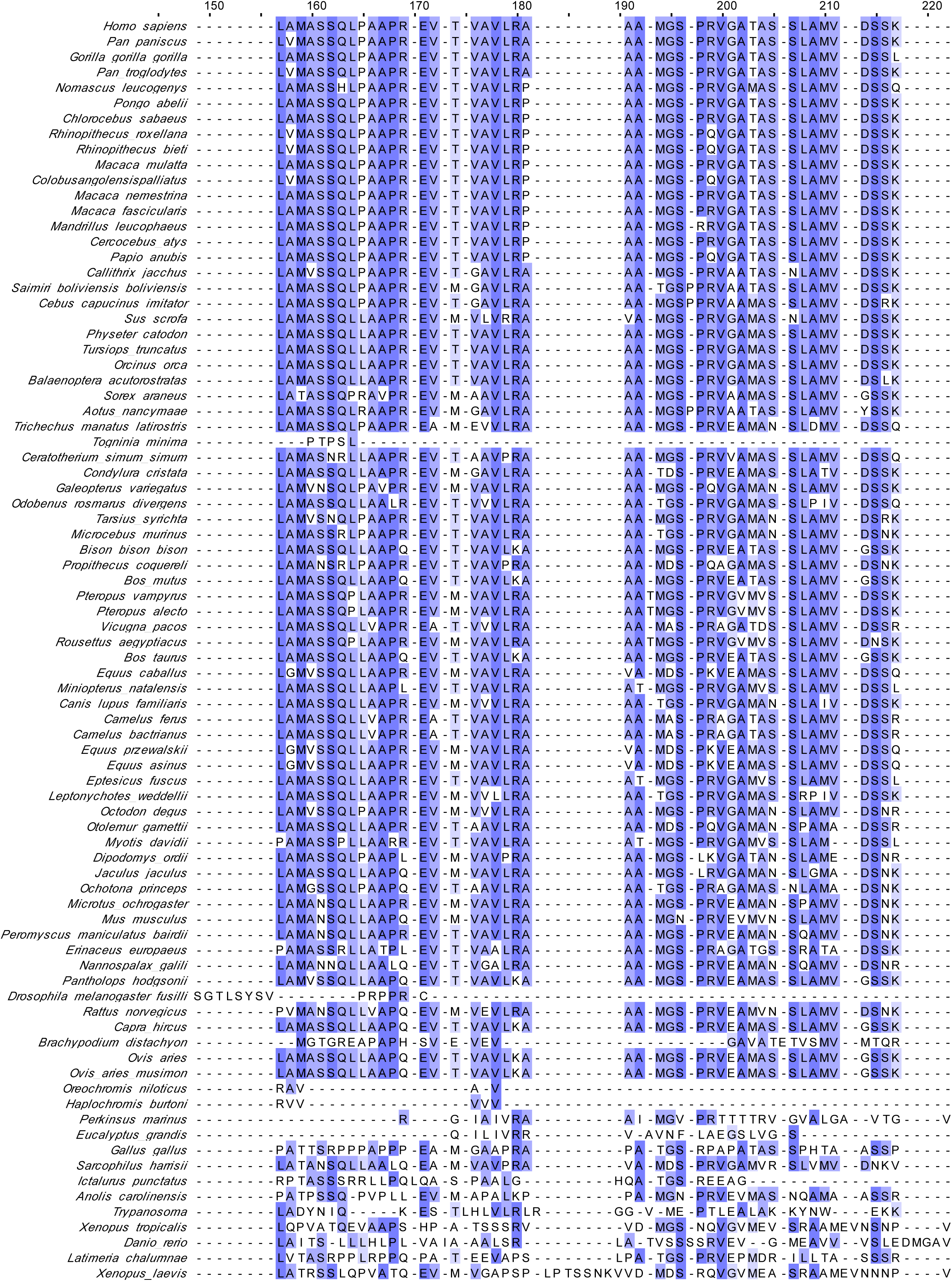
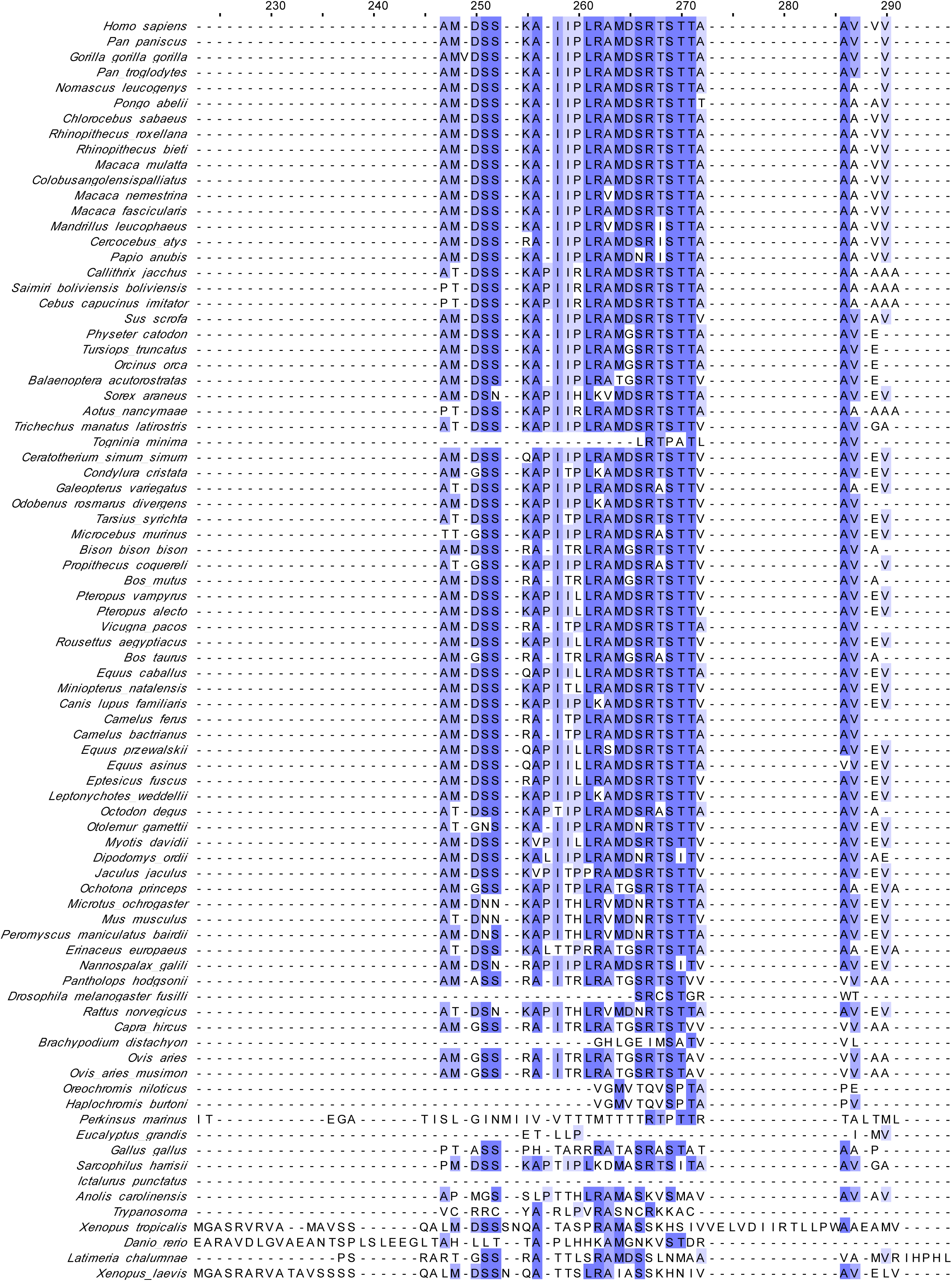
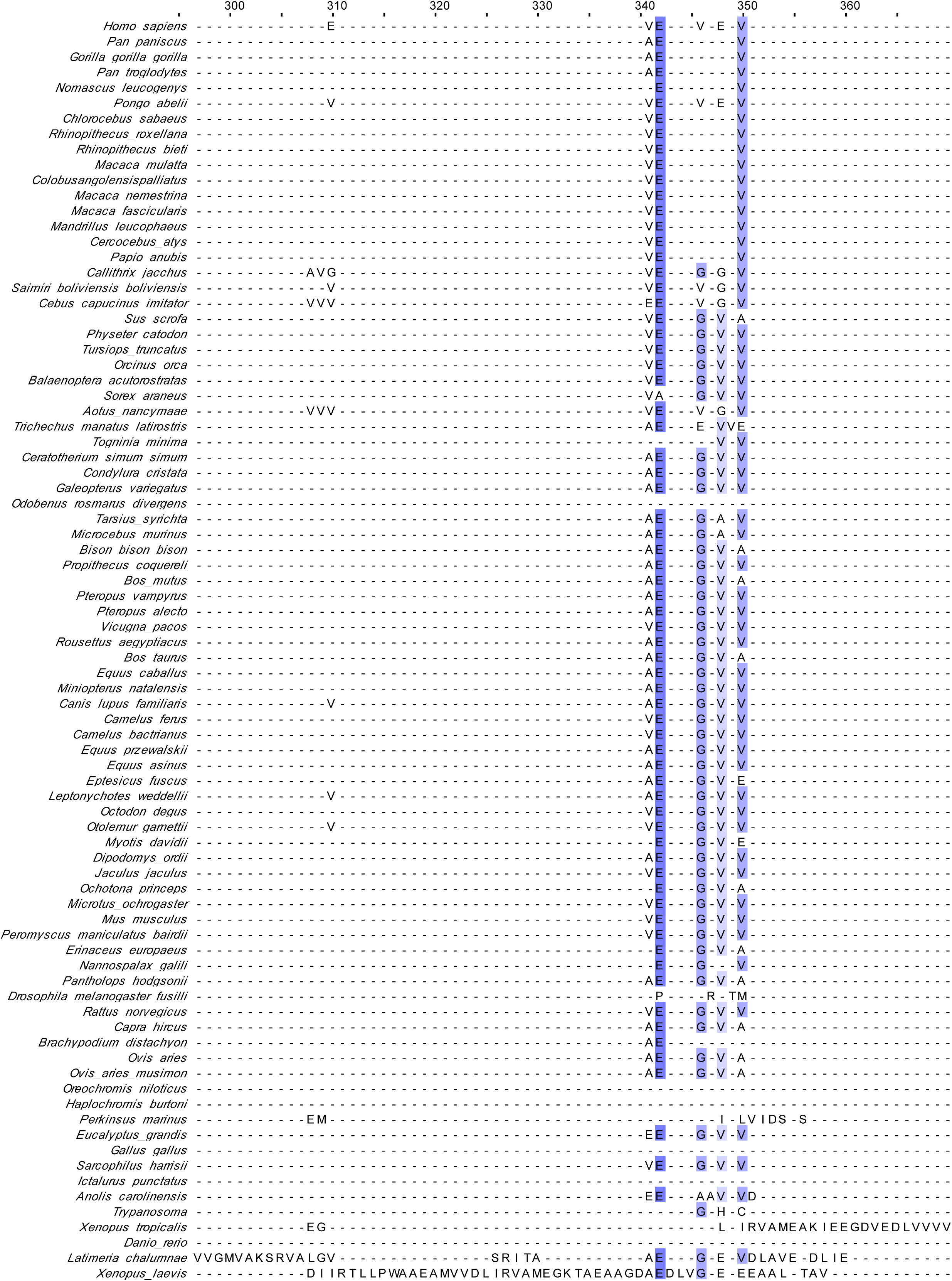
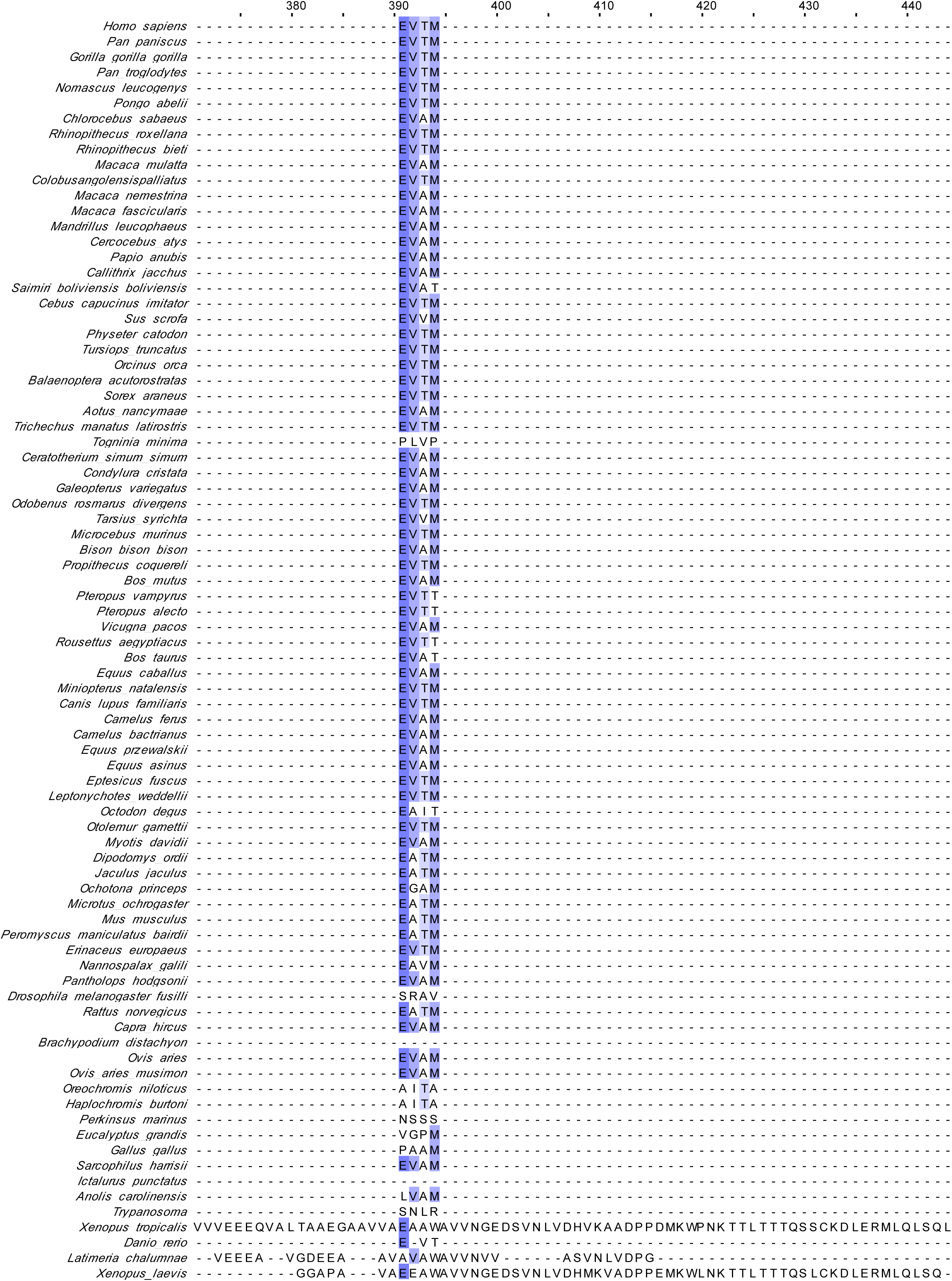
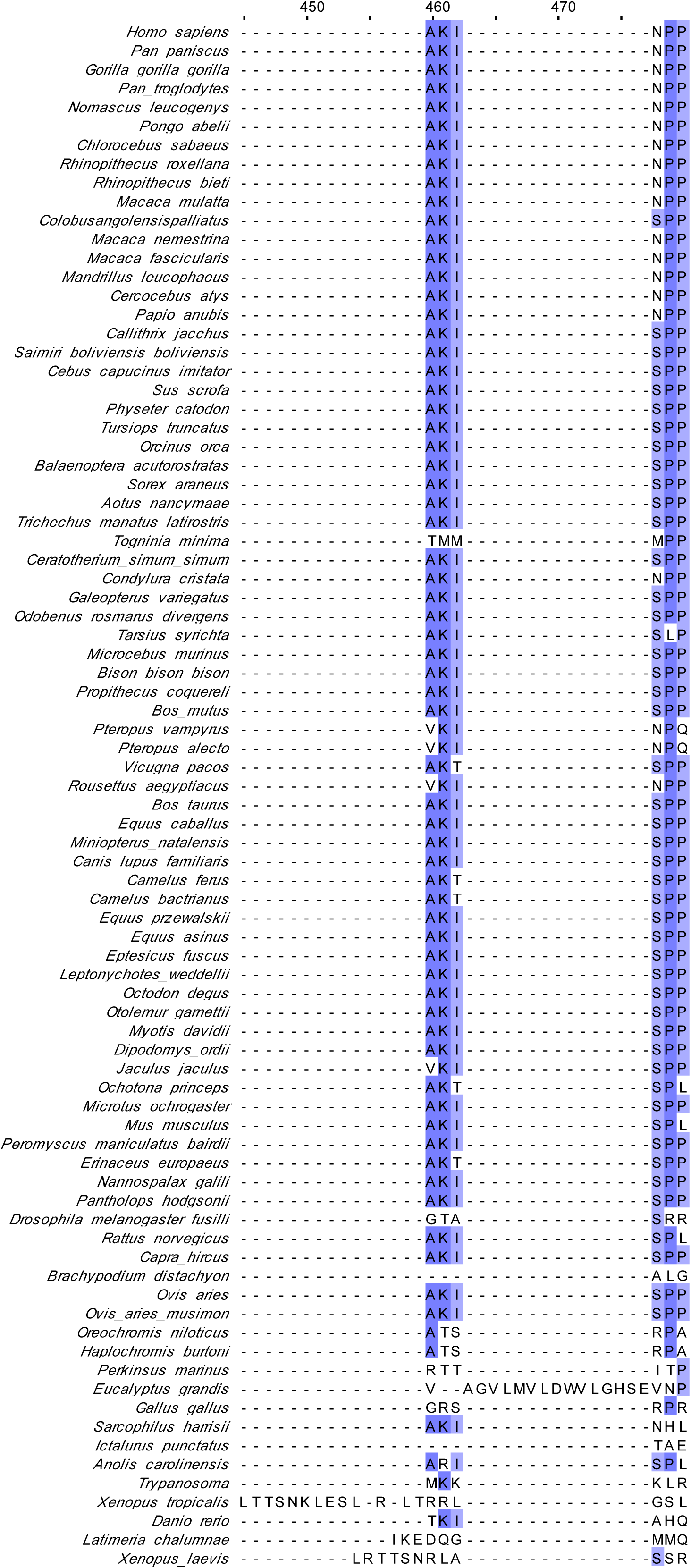
**altFUS protein alignment across species** Alignment of altFUS protein sequences across all 82 species. The alignment was done using Clustalω, and coloured based on conservation from white (highly variable residue) to dark blue (highly conserved residue).

